# A small molecule antagonizes jasmonic acid perception and auxin responses in vascular and non-vascular plants

**DOI:** 10.1101/2021.02.02.429350

**Authors:** Andrea Chini, Isabel Monte, Gemma Fernández-Barbero, Marta Boter, Glenn Hicks, Natasha Raikhel, Roberto Solano

## Abstract

The phytohormone JA-Ile regulates many stress responses and developmental processes in plants. A co-receptor complex formed by the F-box protein COI1 (Coronatine Insensitive 1) and a JAZ (Jasmonate ZIM-domain) repressor perceives the hormone. JA-Ile antagonists are invaluable tools for exploring the role of JA-Ile in specific tissues and developmental stages, and for identifying regulatory processes of the signalling pathway. Using two complementary chemical screens, we identified three compounds that exhibit a robust inhibitory effect on both the hormone-mediated COI-JAZ interaction and degradation of JAZ1 and JAZ9 *in vivo*. One molecule, J4, also restrains specific JA-induced physiological responses in different angiosperm plants, including JA-mediated gene expression, growth inhibition, chlorophyll degradation and anthocyanin accumulation. Interaction experiments with purified proteins indicate that J4 directly interferes with the formation of the Arabidopsis (*Arabidopsis thaliana)* COI1-JAZ complex otherwise induced by JA. The antagonistic effect of J4 on COI1- JAZ also occurs in the liverwort *Marchantia polymorpha*, suggesting the mode of action is conserved in land plants. Besides JA signalling, J4 works as an antagonist of the closely-related auxin signalling pathway, preventing TIR1/Aux-IAA interaction and auxin responses *in planta,* including hormone-mediated degradation of an auxin repressor, gene expression and gravitropic response. However, J4 does not affect other hormonal pathways. Altogether, our results show that this dual antagonist competes with JA-Ile and auxin, preventing the formation of phylogenetically related receptor complexes. J4 may be a useful tool to dissect both the JA-Ile and auxin pathways in particular tissues and developmental stages since it reversibly inhibits these pathways.

**One sentence summary:** A chemical screen identified a molecule that antagonizes jasmonate perception by directly interfering with receptor complex formation in phylogenetically distant vascular and non-vascular plants.

## Introduction

Plant hormones are bioactive small signalling molecules directly perceived by plant receptor complexes. Phytohormones regulate, alone or in combination, multiple aspects of plant development, responses to the environment and to biotic challenges primarily through transcriptional reprogramming (Smith & Li, 2017). Phytohormones regulate most known plant defences and different plant-interacting organisms have evolved the capability to produce phytohormones or phytohormone mimics to induce disease susceptibility and counteract plant defences (Robert-Seilaniantz *et al*. 2011, Pieterse *et al*. 2012, Fonseca *et al*, 2018). For example, the phytotoxin coronatine (COR) is a JA-Ile (jasmonoyl-L-isoleucine) functional analogue produced by *Pseudomonas syringae* to increase its virulence hijacking the plant defence signalling network (Kloek *et al*. 2001, Brooks *et al*. 2004, Gimenez-Ibanez and Solano 2013). In addition to the evolutionary relevance, natural mimic molecules are also important tools in research; for instance, the use of coronatine was determinant to the identification of several components of the JA signalling cascade including the JA-Ile and COR co-receptor COI1 (Coronatine Insensitive 1) (Xie *et al*. 1998, Sheard *et al*. 2010). In addition, several plant hormone analogues are synthetically synthesized and exploited for agricultural and research purposes. For example, several synthetic auxin analogues and anti-auxins represent “classical” herbicides (Van Overbeek and Velez 1946, Grossmann 2010). Mutants completely impaired in auxin perception are not viable; however, auxin-related molecules, such as auxinole and PEO-IAA, are commonly employed to reversibly block auxin perception and study auxin-regulated processes in several plant species (Hayashi *et al*. 2012, Leyser 2018). Therefore, the identification of new molecules interfering with hormonal signalling cascades may be very useful in agriculture and research.

Chemical genomics aims to identify small molecules modifying different biological processes (Norambuena *et al*. 2009, Toth and van der Hoorn 2010, Hicks and Robert 2014). The structural diversity of small compounds is extraordinarily broad (Dobson 2004). Large chemical libraries are screened to identify molecules affecting the activity of a protein, protein families or a pathway in a spatial-temporal and reversible manner. An established advantage of this methodology is to overcome the common limitations of genetic mutant tools such as redundancy or lethality of essential genes (Toth and van der Hoorn 2010, Hicks and Raikhel 2012). Chemical genomic approaches successfully identified hormone analogues or previously unidentified functions of phytohormones (Hicks and Raikhel 2012). A paradigmatic example is the agonist of abscisic acid (ABA) pyrabactin, originally identified as a synthetic inhibitor of growth and cell expansion (Zhao *et al*. 2007). Use of pyrabactin was instrumental in discovering the redundant ABA receptors PYR/PYL (Park *et al*. 2009). Furthermore, chemical genomics helped to uncover the function of strigolactone in light response (Tsuchiya *et al*. 2010).

Jasmonates are lipid-derived phytohormones that mediate responses to abiotic and biotic stress such as drought, salinity, wounding and pathogen attacks (Kazan 2015, Ebel *et al*. 2018, Wasternack and Feussner 2018). In addition, jasmonates regulate the biosynthesis of many secondary metabolites often involved in plant defence, such as anthocyanins or terpenoids (Howe *et al*. 2018, Wasternack and Feussner 2018). Jasmonates are also involved in many developmental processes including growth inhibition and senescence (Howe *et al*. 2018, Wasternack and Feussner 2018). Endogenous developmental processes as well as plant adaptation to environment induce the accumulation of the bioactive form of the hormone, (+)-7-*iso*-JA-Ile (Fonseca *et al*. 2009, Sheard*, et al.* 2010). The identification of the core JA-Ile pathway components evidenced the striking mechanistic parallelism between the JA-Ile and auxins pathways. Both pathways are composed by three elements, *i*) redundant transcription factors regulating JA-Ile or auxins responses (e.g., MYCs or ARFs (Auxin Responsive Factors), respectively), *ii*) a family of redundant repressors of the TF (i.e. JAZs (JAsmonate ZIM-domain) or Aux/IAA (Auxin/Indole-3-Acetic Acid)), which directly or indirectly (through the NINJA adaptor; Pauwels et al., 2010) recruits the co- repressor TOPLESS, and *iii*) a co-receptor formed by an F-box protein (COI1 for JA-Ile and TIR1(Transport Inhibitor Response1)/AFBs (Auxin F-Box) for auxin) and its repressors targets (JAZs for JA-Ile and Aux/IAA for auxins; Chini et al. 2016; Leyser, 2018). In accordance to the “molecular glue” model, in both cases the hormone binding to its co-receptor triggers ubiquitination and degradation of the repressor and, therefore, releases the corresponding TF to activate the pathway (Chini*, et al.* 2007, Maor *et al*. 2007, Thines*, et al.* 2007, Chini *et al*. 2009a, Saracco *et al*. 2009, Lumba *et al*. 2010, Leyser 2018, Chini *et al*. 2016, Howe*, et al.* 2018) . It is remarkable that the receptors COI1 and TIR1/AFBs have a common evolutionary origin and, although strikingly similar, they are adapted to perceive structurally different ligands (Bowman *et al*. 2017).

The JAZ proteins are repressors that inhibit a large number of TFs belonging to several different families, including bHLH (basic helix-loop-helix), MYB, and EIN3/EIL (Chini*, et al.* 2016, Howe*, et al.* 2018, Zander *et al*. 2020). The identification of these components of the JA perception and signalling pathway were carried out using mutant screens or reverse genetics (Xie*, et al.* 1998, Browse 2009, Chini*, et al.* 2016). However, several components of the JA-signalling pathway belong to large protein families with redundant functions, such as the 13 Arabidopsis JAZ repressors or the 4 MYC transcription activators, where genetic approaches are very laborious and their success may be more limited (Chini*, et al.* 2007, Thines*, et al.* 2007, Fernandez-Calvo *et al*. 2011, Qi *et al*. 2015, Guo *et al*. 2018, Zander *et al*. 2020). Therefore, specific antagonist molecules of JA perception would represent excellent tools to overcome redundancy and to explore the role of JA in spatiotemporal analyses. A chemical screen identified jarin1 as inhibitor of jasmonate responses, impairing the synthesis of JA-Ile (Meesters *et al*. 2014). In addition, coronatine-*O*-methyloxime (COR-MO) was rationally designed as specific antagonist of jasmonate perception that shows strong inhibiting activity of COI1-JAZ interaction, JAZ degradation and several JA-mediated responses (Monte *et al*. 2014).

In this work, we searched for different and cheaper antagonists of JA-Ile perception by chemical screens. Here we describe the identification of three commercially available JA-Ile antagonists and their characterization. These molecules prevent the JA-Ile-mediated COI-JAZ interaction and the degradation of JAZ1 and JAZ9 *in vivo*. Moreover, one molecule (J4) exhibited JA-antagonist effects *in planta*, preventing several JA-mediated responses such as gene expression, growth inhibition, chlorophyll degradation and anthocyanin accumulation in Arabidopsis, tomato (*Solanum licopersicum*) and *Nicotiana benthamiana*. In addition to JA signalling, J4 also inhibited the closely-related auxin pathway but no other hormonal pathways. Furthermore, the mode of action of this molecule as direct antagonist of JA-Ile perception by COI1-JAZ complexes is conserved in land plants, indicating its potential use in any plant species. This commercially available compound is a potential powerful tool for the pharmacological analysis and dissection of the JA and auxin signalling pathways.

## Results

### Chemical Screens

The perception of the bioactive form of the JA-Ile hormone is mediated by the co-receptor COI1-JAZ (Sheard*, et al.* 2010). In order to identify synthetic molecules promoting or interfering with the COI1-JAZ interaction, we carried out two independent chemical screens based on the interaction between COI1 and JAZ9 in yeast-two-hybrid assays (Chini *et al*. 2009b, Fonseca*, et al.* 2009). Three chemical libraries containing approximately 22,500 molecules (see Material and Method) were tested to identify agonists and antagonists of JA-Ile perception, inducing or preventing the hormone-triggered COI1-JAZ9 interaction, respectively. None of the agonist compounds found could reproducibly induce COI1-JAZ9 interaction. However, 5 antagonist molecules were identified and confirmed (Y10, Y11 Y17, Y18 and Y20; Y defines molecules identified in the Y2H screen, Figure 1A and B, Suppl. Figure S1A and Supplemental Table S1), with Y10 being weaker than the other four. We also confirmed that these compounds do not affect other interactions of these proteins (JAZ9 with itself or with NINJA) ruling out general unspecificity. We also tested whether these molecules could inhibit the formation of additional COI1-JAZ complexes, confirming that these five compounds prevent the COR-induced COI1-JAZ3 interaction (Suppl. Figure S1B).

**Figure 1.**
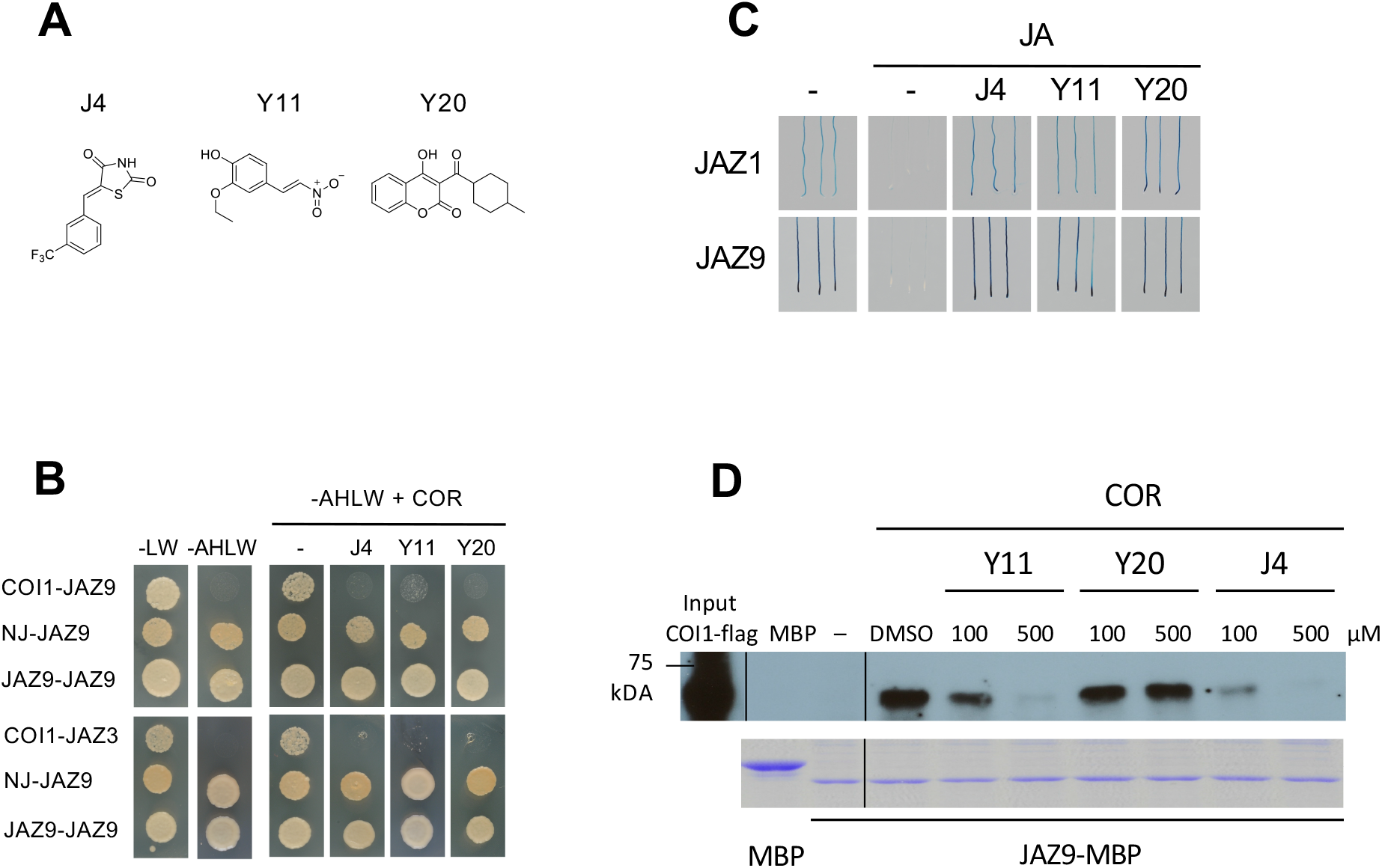
**Identification of molecules with JA-Ile antagonist activity in yeast two- hybrid, *in planta* and in the hormone-dependent formation of the receptor complex COI1-JAZ9.** (A) Chemical structure of the JA-antagonist compounds. (B) Yeast cells co-transformed with pGAD-JAZ9 (preys) and pGBK-COI1, pGBK- NINJA or pGBK-JAZ9 (baits) were selected and subsequently grown on yeast synthetic drop-out lacking Leu and Trp (-LW) as a transformation control, or on selective media lacking Ade, His, Leu and Trp (-AHLW) to test protein interactions. COI1 interaction with JAZ9 or JAZ3 is detected only in presence 5 mM or 20 mM coronatine respectively. Antagonist molecules inhibit COI1-JAZ interaction in presence of COR (J4 and Y20 were used at 15 μM, and Y11 at 300 nM). Compounds were dissolved in DMSO; therefore, an equivalent volume of DMSO was used in the negative control (labelled as -). As control, we tested that the antagonist molecules did not interfere with the interaction between JAZ9 and NINJA or JAZ9 dimerization. (C) 6-day-old JAZ1-GUS and JAZ9-GUS seedlings were concurrently treated with 2.5 μM JA and the indicated molecules (J4 and Y20 were used at 25 μM, and Y11 at 50 μM). Jasmonic acid triggers the degradation of JAZ-GUS protein, whereas the addition of antagonist molecules prevented JAZ degradation. Compounds were dissolved in DMSO; a volume of DMSO equivalent to that in the compounds was used in the negative control (identified as -). (D) Immunoblot analysis of COI1-flag/JAZ9-MBP interaction performed with anti-flag antibodies to COI1-flag protein recovered from MBP-JAZ9 and extracts of transgenic *COI1-flag* plants. MBP was employed as negative control. The COI1-flag/JAZ9-MBP interaction is dependent on the presence of COR (0.5 μM). Compounds were dissolved in DMSO; therefore, a volume of DMSO equal to that in 0.5 μM of the compounds was used as control. The compounds Y11 and J4 reduce COI1-flag/JAZ9-MBP interaction (at 100 and 500 μM). The lower panel shows Coomassie blue staining of the MBP and the JAZ9-MBP after cleavage with Factor Xa.

Two complementary chemical screens were also carried out *in planta* exploiting the rapid hormone-dependent JAZ degradation using the 35S:*JAZ1*-GUS reporter line (Thines*, et al.* 2007). Approximately 15,000 compounds (see Material and Methods) were tested for triggering degradation of JAZ1-GUS in the absence of JA-Ile/COR or preventing COR-induced degradation of JAZ1-GUS. Again, no agonist compounds were confirmed, whereas we validated 7 antagonist molecules (J1, J2, J3 J4, J9, J10 and J11; J defines the compounds identified in the JAZ1-GUS screen), preventing hormone-induced degradation of JAZ1 (Figure 1A and C, Suppl. Figure S2 and Supplemental Table S1). Since the original screen was carried out with COR, we tested the candidate molecules in response JA to discard unspecific effects of COR or an effect of the compound on JA conversion into JA-Ile. All J1 to J11 compounds were confirmed (Suppl. Figure S2).

Among the 7 compounds obtained in the screens *in planta* (J) only one was also active in yeast assays, whereas among the 5 compounds identified in Y2H screens (Y) only 3 were positive *in planta* (Suppl. Figure S3; despite Y10 was positive we discarded this compound because of its weak activity both in yeast and plant). Therefore, we confirmed 3 compounds (J4, Y11 and Y20; Figure 1A) with robust antagonistic activity in both Y2H and *in vivo* assays. These three compounds prevented the JA-triggered degradation of JAZ1-GUS and JAZ9-GUS *in vivo*, showing that their effect is not restricted to a particular JAZ (Figure 1B). Similarly, the three compounds inhibited the interaction of COI1-JAZ9 and COI1-JAZ3 in Y2H, without affecting other interactions of these proteins (JAZ9 with itself or with NINJA, Figure 1C). The effect of J4 and Y11 as antagonists of COR-induced COI1-JAZ9 interaction was confirmed in semi-*in vivo* pull-down (PD) experiments between recombinant JAZ9 fused to maltose binding protein (MBP) and COI1-flag expressed *in planta* (Figure 1D).

In summary, these results confirmed that two molecules, namely J4 and Y11, interfere with the COI1 interaction with more than one JAZ co-receptor both in the heterologous yeast system and in PD experiments. This suggests that these compounds might compete with COR or JA-Ile for the binding to COI1 *in vivo*. However, an indirect effect of these molecules acting through a regulator of COI1 or JAZ cannot be discarded yet.

### Minimal active concentration of antagonist molecules of JA-Ile/COR perception

Next, we estimated the minimal active concentration at which these molecules could act as antagonist of the JA-Ile/COR perception. The J4 and Y20 compounds prevent the formation of the COI1-JAZ receptor complex in yeast at a concentration between 5 and 50 μM, similar to the concentration of COR (5 μM) used to induce COI1- JAZ interaction (Suppl. Figure S4). The compound Y11 inhibits the COI1-JAZ9 interaction at a concentration as low as 300 nM; however, Y11 is toxic for the yeast at a concentration of 1 μM as shown by the lack of yeast growth in the control JAZ9-JAZ9 interaction assay (Suppl. Figure S4).

Similarly, the J4 and Y20 molecules prevent JAZ1-degradation *in planta* at minimal concentrations between 10 and 25 μM, whereas Y11 had a weaker effect (Suppl. Figure S5). In summary, J4 and Y20 show an antagonistic activity on processes mediated by JA-Ile and COR at a concentration close to that usually employed for exogenous hormone treatments.

### Structure-activity relationship

To define the active moiety of these molecules, we performed an analysis of structure-activity relationship of the three identified compounds (Rosado *et al*. 2011). No structurally related compounds of Y11 were available. In contrast, we found three derivatives of Y20 (named Y20-L1, Y20-L2 and Y20-L3, Suppl. Figure S6A). Y20-L1 and Y20-L2 prevented COI1 interaction with both JAZ9 and JAZ3 at the same concentration as Y20 in Y2H (Suppl. Figure S6B). Similarly, they prevented the JA-induced degradation of both JAZ1-GUS and JAZ9-GUS *in planta* (Suppl. Figure S6C), whereas Y20-L3 fails to show antagonistic activity in all bioassays (Suppl. Figure S6B and 6C). These results suggest that the antagonistic activity of Y20 resides in the minimal 3-butyryl-4-hydroxy-2H-chromen-2-one structure. Moreover, we identified that either or both the hydroxy group and the butyryl lateral chain are essential for the activity of this molecule as JA antagonist, as simultaneous shortening of two carbons in the butyryl chain and loss of the hydroxy group in Y20-L3 abolish the activity (Suppl. Figure S6A).

The use of J4 derivatives (Suppl. Figure S6A) showed that the different substitutions of the benzene ring did not have any impact on the activity of these molecules, since all of them retained the activity as JA antagonists *in planta* in the JAZ1 and JAZ9 degradation assays (Suppl. Figure S6C). We concluded that the 5- (benzylidene)-1,3-thiazolidine-2,4-dione moiety is important for the antagonist activity.

### Molecules inhibiting JA-Ile perception affect JA-mediated transcription in vivo

We subsequently analysed the effect of these molecules on JA-induced gene expression *in planta*. *JAZ2* and *JAZ9* are among the genes most rapidly and strongly induced by JA in a COI1-dependent manner (Chini*, et al.* 2007, Thines*, et al.* 2007). Therefore, we assessed the effect of these molecules on the JA-induced gene expression in the *pJAZ2:GUS* and *pJAZ9:GUS* reporter lines (Monte*, et al.* 2014, Gimenez-Ibanez *et al*. 2017). In basal conditions, *pJAZ2:GUS* and *pJAZ9:GUS* are expressed at very low levels but they are quickly and strongly induced in response to JA and after wounding stress, which stimulates the synthesis of endogenous JA-Ile (Figure 2A and 2B; (Monte*, et al.* 2014)). Simultaneous treatment of exogenous JA or wounding with the identified molecules, and derivative compounds, strongly inhibited the JA-mediated activation of both *JAZ2* and *JAZ9* expression *in planta* (Figure 2A-B and Suppl. Figure S7).

**Figure 2.**
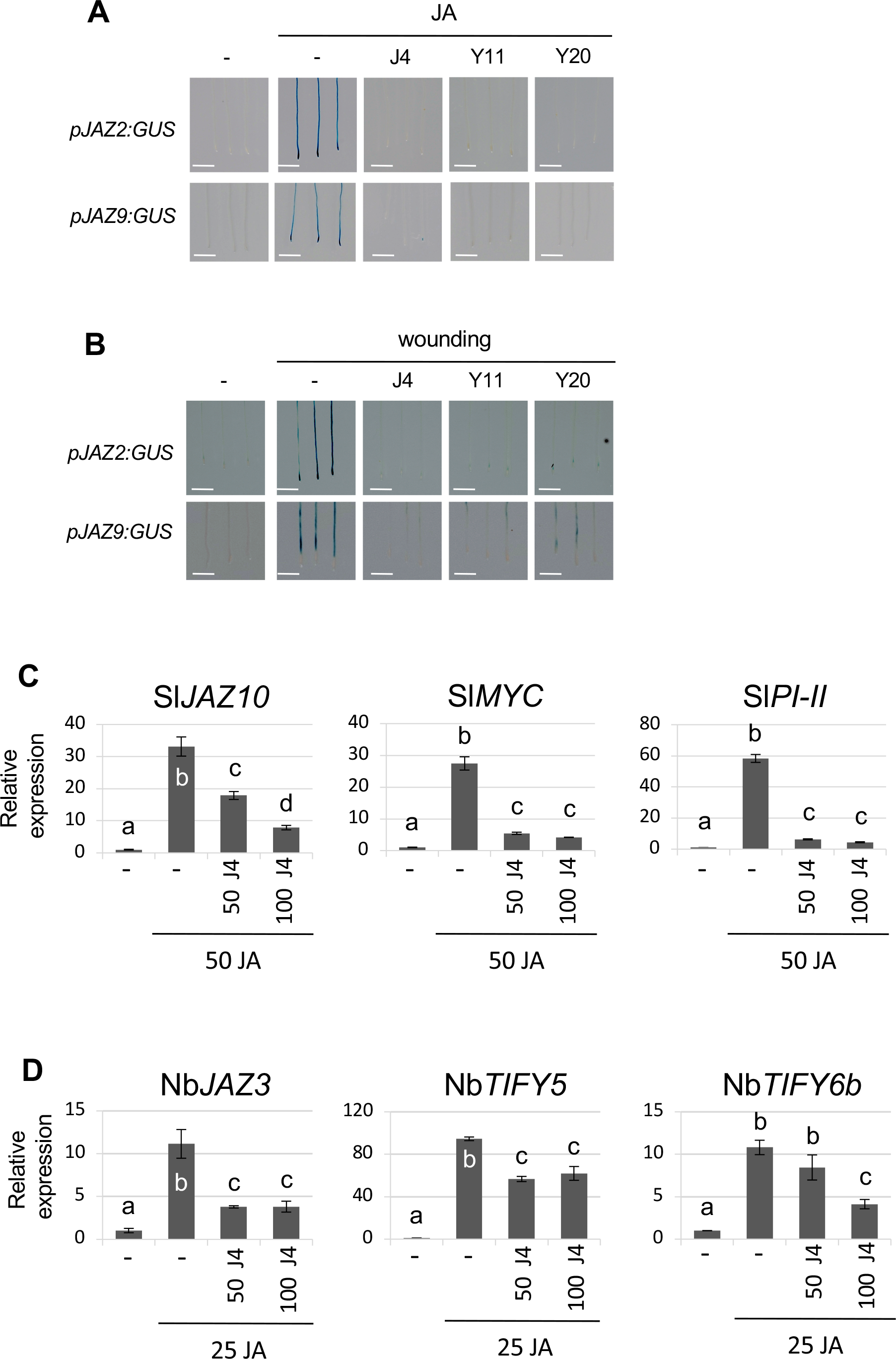
**Antagonistic molecules prevent the JA-induced expression of several *JAZ* genes *in planta*.** (A) Representative seedlings (N > 30) of the *pJAZ2:GUS* and *pJAZ9:GUS* line were concurrently treated with 5 μM JA for 75 minutes and the indicated molecules (J4 at 10 μM, Y20 at 25 μM, and Y11 at 100 μM) for 3 hours. Jasmonic acid triggers the expression of *JAZ*, whereas the addition of antagonist molecules could prevent *JAZ* transcriptional activation. Compounds were dissolved in DMSO; an equivalent volume of DMSO was used in the negative control (identified as -) in A and B. White bars are equal to 1 mm in A and B. (B) Seedlings of *pJAZ2:GUS* and *pJAZ9:GUS* were wounded several times and concurrently treated with the indicated compounds (J4 at 10 μM, Y20 at 25 μM, and Y11 at 100 μM) for 2 hours. The mechanical wounding induces the *JAZ* expression whereas the JA-antagonists can prevent wound-induced *JAZ* expression. (C-D) Gene expression analysis of JA-marker genes in *Solanum lycopersicum*, moneymaker cultivar, (C) and *Nicotiana benthamiana* (D) plants in response to JA treatment and/or the J4 molecule. Tomato (C) plants were pre-treated with DMSO (-; untreated control), 50 or 100 µM J4 for 1 hour and then with 50 µM JA for 1 hour. Sl*αTUB4* was used as housekeeping control gene. *Nicotiana benthamiana* plants were pre-treated with DMSO (-; untreated control), 50 or 100 µM J4 for 1 hour and then with 25 µM JA for 1 hour. Nb*βACT* was used as housekeeping control gene. Experiments were repeated three times with similar results. One-way ANOVA with post-hoc Tukey HSD Test (p< 0.01) analyses define the significant differences in gene expression. Each biological sample consisted of tissue pooled from 10-15 plants. Data show mean ± SD of three to four technical replicates.

Interestingly, the *JAZ9* transcript accumulates specifically in trichomes in basal conditions (Suppl. Figure S8A). This basal expression is abolished in the *coi1-30* background, which indicates that perception of endogenous JA-Ile is required for the specific *JAZ9* expression in the trichomes (Suppl. Figure S8B). We assessed the inhibitory effect of the identified molecules on the trichome tissue specific expression of *JAZ9*. Interestingly, only the compound J4 repressed the JA-regulated expression of *JAZ9* in the trichomes (Suppl. Figure S8C). To further assess the inhibitory effect of J4 on additional JA-markers, we analysed the expression of the JA-induced *OPR3*, *TAT3* and *JAZ10* genes. Indeed, exogenous treatment with J4 partially prevents the transcriptional activation of these markers by JA (Suppl. Figure S9A).

To test if the JA-inhibitory activity of J4 is conserved across angiosperm plants, we tested the effect of J4 on JA-regulated gene expression in two species of the Asterid clade, tomato and *Nicotiana benthamiana*. Similar to the results in Arabidopsis, exogenous treatment with J4 significantly prevents the JA-induced transcriptional activation of several marker genes in both tomato and *N. benthamiana* plants (Figure 2 C-D).

These results show that these three compounds can prevent transcriptional activation induced by both endogenous and exogenous JA. Moreover, J4 can also inhibit the expression of a marker gene mediated by endogenous JA-Ile in a specific tissue as well as in different angiosperm plants.

### Physiological effect of the identified antagonists in planta

Jasmonate induced several physiological responses, such as growth inhibition, anthocyanin accumulation and reduction of chlorophylls, among others (Wasternack and Feussner 2018). In order to assess the effect of the identified molecules on JA-regulated physiological responses, we grew plants in presence of JA and the identified molecules. The concurrent treatment of the J4 compound at concentrations as low as 5 μM could partially prevent the root growth-inhibition, chlorophyll degradation and anthocyanin accumulation induced *in planta* by JA (10 μM; Figure 3). In contrast, Y11 and Y20 failed to prevent JA-mediated responses *in planta* (Suppl. Figure S10). We also determined the half maximal inhibitory concentration (IC50) of J4. For a concentration of 15 μM of JA in the JAZ1-GUS degradation assay, the IC50 of J4 was 17.66 μM (Suppl. Figure S11). To study if J4 conserved its JA-inhibitory activity in different plants, we tested the effect of J4 on JA-induced chlorophyll and carotenoids degradation in *N. benthamiana*. Exogenous treatment of *N. benthamiana* with J4 significantly prevents the JA-induced degradation of both chlorophyll and carotenoids *in planta* (Suppl. Figure S12).

**Figure 3.**
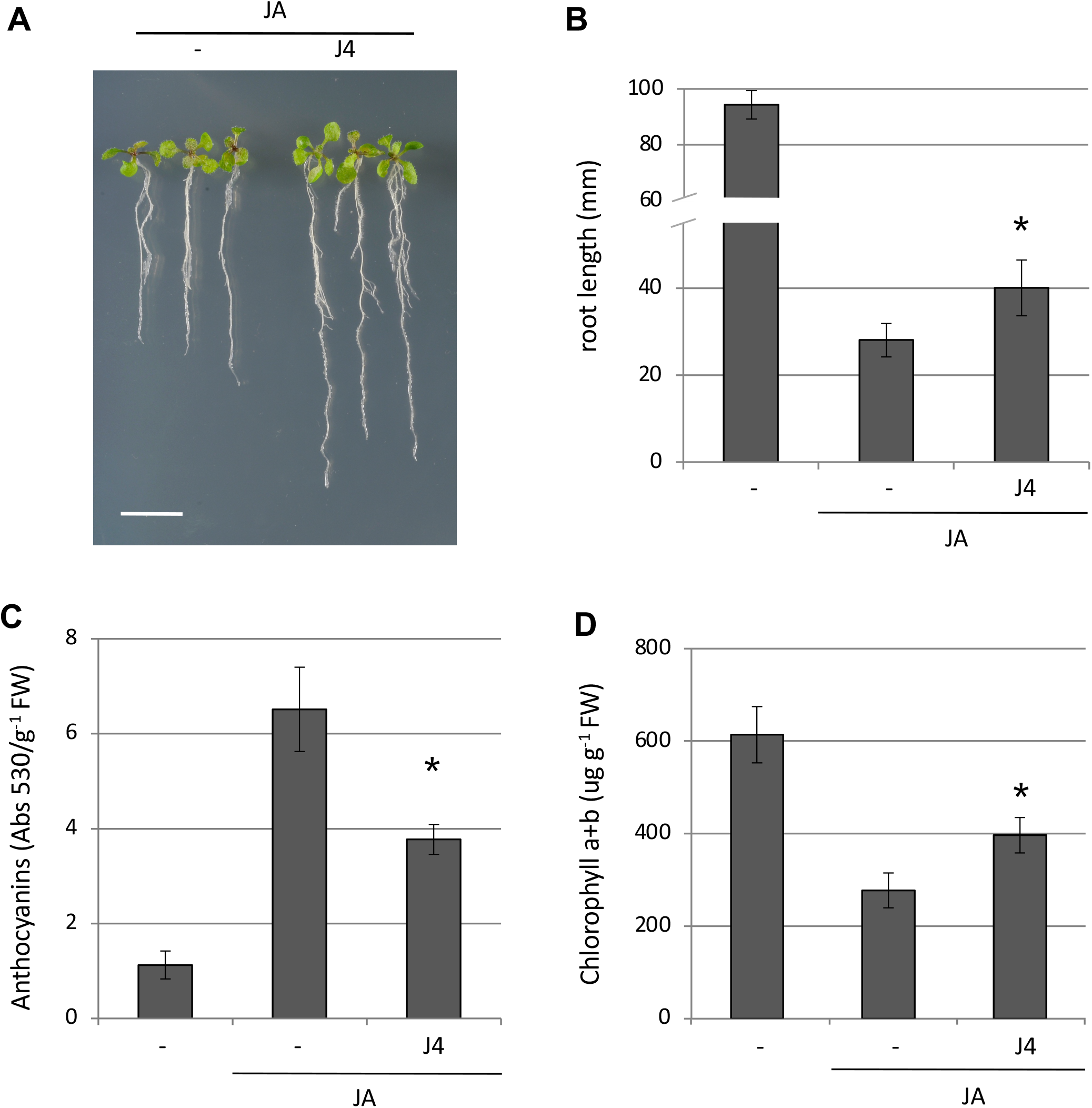
**Antagonistic effect of the compound J4 on JA-induced responses *in planta*.** (A) 13-day-old WT seedlings grown for 10 days on vertical plates in presence of 10 μM JA with or without 5 μM J4. White bar stands for 1 cm. (B) Root growth inhibition by 10 μM JA of 13-day-old seedlings in presence or absence of 5 μM J4. J4 was dissolved in DMSO; therefore, an equivalent volume of DMSO was used in the negative control (identified as -) in A-D. (C) Anthocyanin accumulation, shown as Absorbance (530) per gram of plant fresh weigh (FW), in 7-day-old WT seedlings grown for 2 days in presence of 10 μM JA with or without 2.5 μM J4. (D) chlorophyll a and b content of 13-day-old WT seedlings grown for 10 days on vertical plates in presence of 10 μM JA with or without 5 μM J4. (B, C and D) Bars represent the average value and error bars the standard deviation. Asterisks indicate statistically significant values of JA-treated plants according to a Student’s t-test (p < 0.05).

Altogether, these data confirm that J4 acts as an antagonist of the JA pathway and inhibits several hallmark JA-mediated responses *in planta*.

### Specificity of J4

To study J4 specificity, we analyzed possible inhibitory side effects of J4 on the ubiquitin/proteasome process that would in turn stabilize the JAZ proteins. Therefore, we analysed the effect of J4 on other hormonal pathways regulated by the ubiquitin/proteasome system, similarly to the JA pathway. First, we monitored the gibberellin-mediated proteasome degradation of RGA-GFP (Repressor of GA) and the constitutive proteasome-dependent degradation of EIN3-GFP (Ethylene-Insensitive-3), which is inhibited by ethylene or its precursor ACC (Silverstone *et al*. 2001, Guo and Ecker 2003). As shown in Figure 4A, J4 did not prevent the GA-triggered degradation of RGA. Similarly, J4 did not substantially inhibit the constitutive degradation of EIN3 in the absence of ACC (Figure 4B). Secondly, we assessed the effects of J4 on the turnover of the auxin Aux/IAA repressors, whose degradation depends on the activity of the F-box TIR1, the receptor of auxins and the closest homolog of COI1 (Chico *et al*. 2008). In contrast to GA and ethylene, J4 partially inhibited the auxin-mediated degradation of the dII-VENUS auxin-repressor marker and the auxin-repressor IAA1 (Figure 4C and 4D). To further assess the effect of J4 on the auxin-pathway, we tested the impact of this molecule on the auxin transcriptional marker *DR5:GUS*. Exogenous indole-3-acetic acid (IAA) treatment highly induced the expression of *DR5:GUS*, whereas the anti-auxin TIBA (2,3,5-triiodobenzoic acid) and J4 could partially inhibit the IAA-mediated *DR5:GUS* expression (Figure 5A). PD analyses in Fig 5B shows that IAA induces the interaction between TIR1 and IAA7 whereas J4 partially inhibited this interaction, similarly to the effect of the specific auxin perception inhibitor auxinole (Figure 5B). Finally, J4 also impairs the auxin-regulated gravitropic response in root (Figure 5C-D).

**Figure 4.**
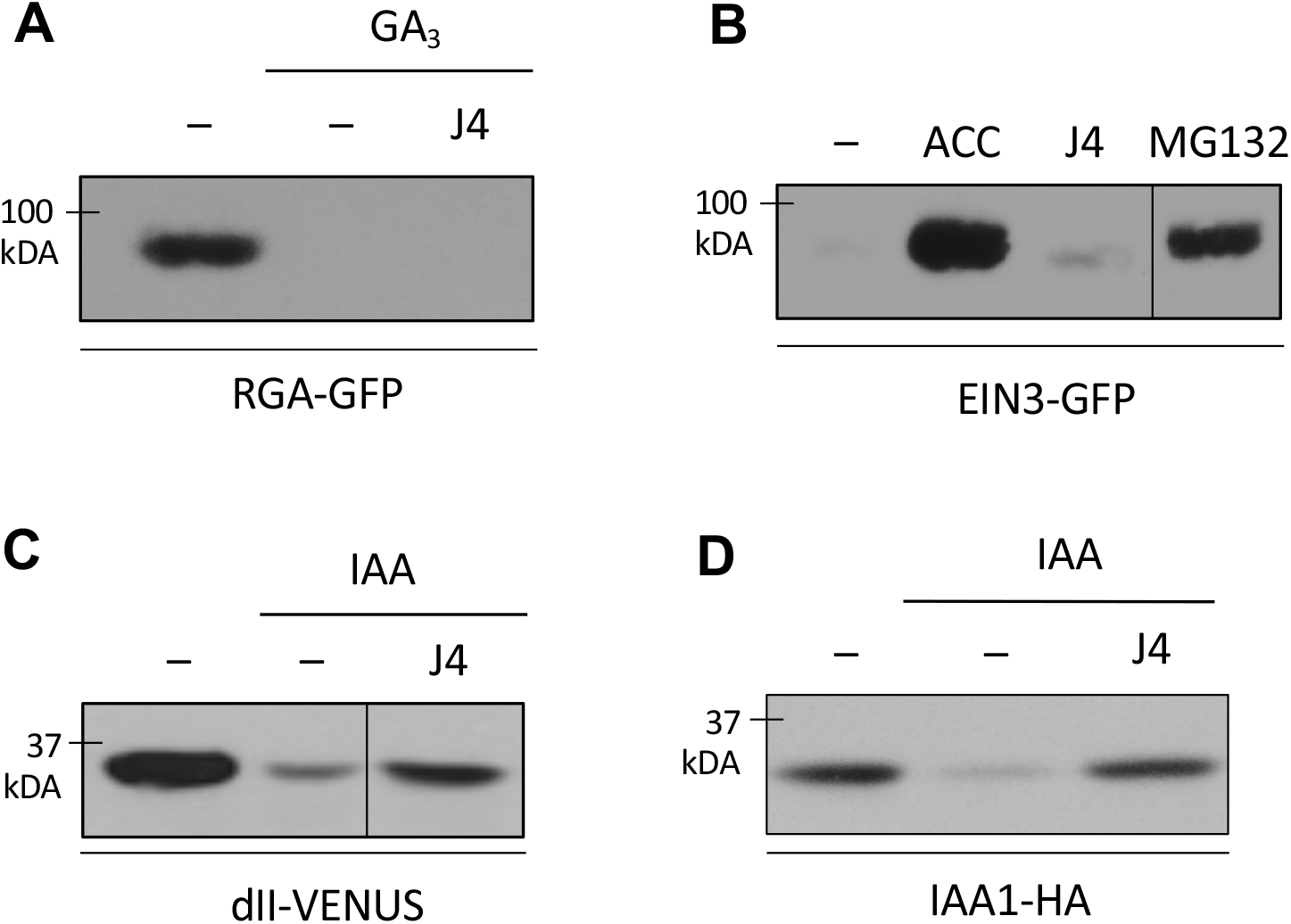
**The compound J4 does not affect the ubiquitin/proteasome system.** (A) Immunoblot analysis of the effect of antagonists on RGA-GFP (Repressor of GA) stability (with anti GFP antibodies). pRGA:GFP-RGA seedlings were concurrently treated for 2 hours with 10 µM gibberellic acid (GA_3_) and J4 at 10 µM. GA_3_ induces RGA degradation and J4 fails to prevent RGA destabilization. J4 was dissolved in DMSO; an equivalent volume of DMSO was used in the negative control (labelled as -) in A-D. (B) Immunoblot analysis of the effect of antagonists on EIN3-GFP (Ethylene- Insensitive-3) stability (with anti GFP antibodies). Seedlings of the 35S:EIN3-GFP line were treated for 2 hours with the J4 at 10 µM or 10 µM of the ethylene precursor ACC. EIN3 is continuously degraded and stabilized by ACC. The synthetic proteasome inhibitor MG132 (at 100 µM) was included as positive control. (C-D) Effect of J4 the stability of the on dII-VENUS (with anti GFP antibodies) auxin sensor and IAA1-HA (with anti HA antibodies) *in planta*. Immunoblot analysis of 35S:dII-VENUS and 35S:IAA1-HA seedlings of the dII-VENUS auxin repressor marker (C) or the auxin repressor IAA1 (D) treated for 1 hour with 1 µM IAA in absence or presence of J4 at 10 µM. The first band shows constitutive expression of the dII-VENUS auxin marker (C) or IAA1-HA (D).

**Figure 5.**
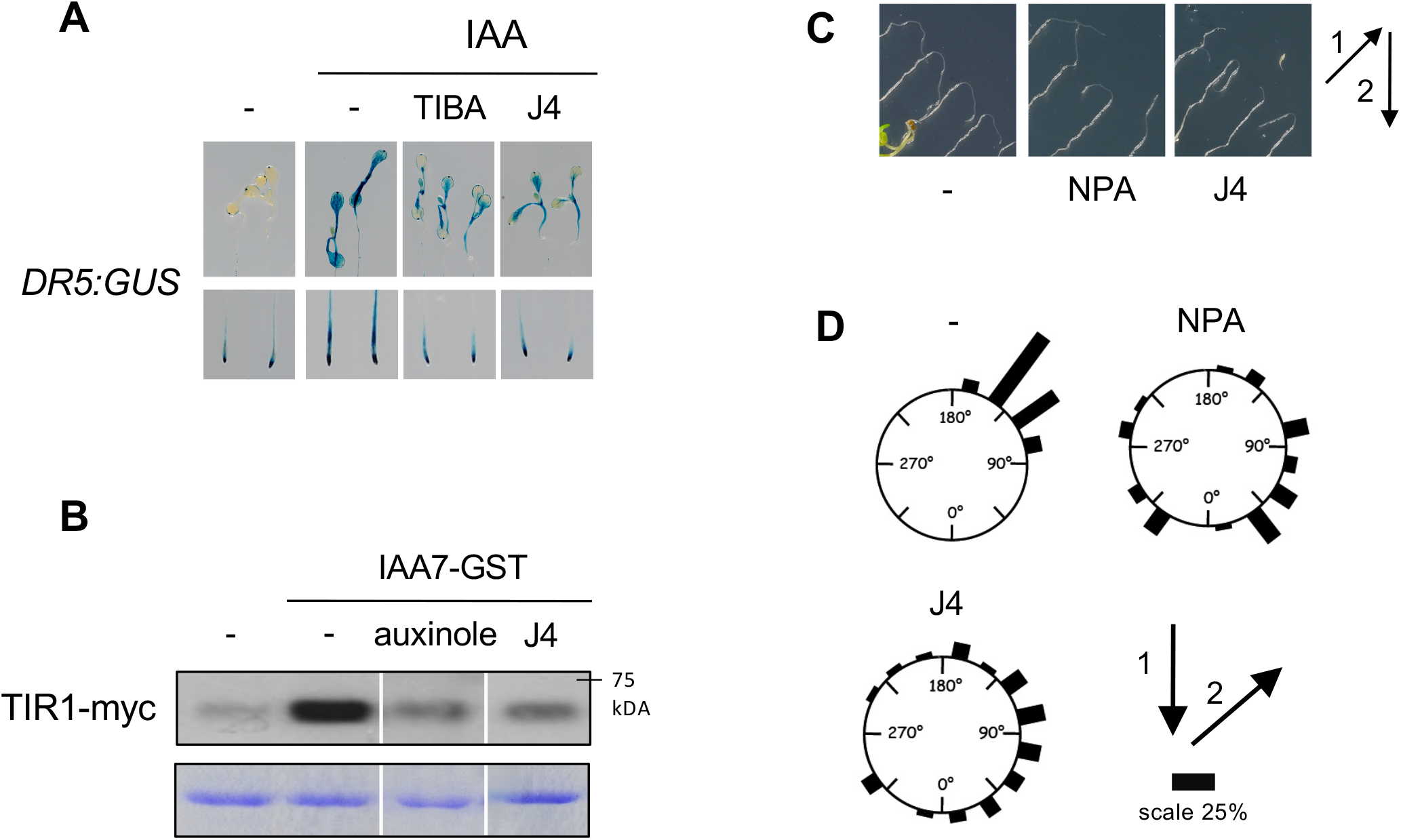
**Effect of the compound J4 on several auxin-regulated responses.** (A) Effect of J4 on auxin-induced *DR5:GUS* reporter gene expression. 6-d-old seedlings were treated with IAA (5 μM) alone and in presence of the auxin inhibitor TIBA (100 μM) or J4 (10 μM) for 90 minutes. J4 was dissolved in DMSO, therefore an equivalent volume of DMSO was used as negative control (defined as -) in A-D. (B) Immunoblot analysis of TIR1-myc/IAA7-GST interaction performed with anti-myc antibodies to TIR1-myc protein recovered from IAA7-GST and extracts of transgenic *TIR1-myc* plants. IAA (5 μM) induces the interaction between the auxin co-receptors TIR1-myc and IAA7-GST, whereas the specific auxin-perception inhibitor auxinole and J4 (at 100 μM) partially inhibits the TIR1-IAA7 complex formation. (C-D) Effect of J4 on auxin-mediated gravitropic response. 4-day-old seedlings grown on MS plates were transferred to control DMSO ( - ), NPA (1 μM) or J4 (2.5 μM) plates and then rotated 135° from the gravity vector (arrow 1 vs 2). Photographs were taken 24 hours after re-orientation (C). Bar lengths represent the percentage of seedlings (n = 28 to 32 seedlings for each treatment) growing at the indicated orientation 24 hours after re-orientation (D).

These results show that J4 does not generally affect ubiquitin/proteasome-regulated systems (UPS) but is not a fully specific antagonist of JA-Ile perception, having also an inhibitory effect on the auxin pathway.

### J4 mode of action: A direct antagonist of hormone receptor complex

Previous results suggest a possible mode of action of J4 as a direct competitor with JA-Ile or COR for induction of COI1-JAZ complex (or competition with auxin in the case of TIR1-IAA complex). To further study the J4 mode of action on the JA-pathway, we checked if this molecule interferes directly with the assembly of the COI1- JAZ perception complex *in vitro*, in the absence of other plant proteins. For that, we used a heterologous baculovirus–insect cell expression system to express and purify the COI1-flag protein. Since purified COI1 is quite unstable and mostly inactive, we also expressed in insect cells a key component of the SCF^COI1^ complex, the adaptor Arabidopsis SKP-like protein 1 (ASK1), which stabilizes and maintains the bioactivity of COI1 (Li *et al*. 2017). In the presence of ASK1, COR promotes the interaction between the purified COI1 and several recombinant JAZ-MBP proteins (JAZ1, JAZ2, JAZ3 and JAZ9). J4 efficiently inhibits the COR-induced formation of all the tested COI1-JAZ complexes (Figure 6A). To test if the mode of action of J4 is conserved across land plants, we tested the effect of J4 on the MpCOI1-MpJAZ co-receptor from the bryophyte *Marchantia polymorpha* (Monte *et al*. 2018). This co-receptor is the most different COI1-JAZ complex known from that of Arabidopsis, which even perceives a different ligand (dn-OPDA instead of JA-Ile), and diverged evolutionarily more than 450 million years ago (Bowman *et al*. 2017; Monte *et al*. 2018). The MpCOI1 and MpASK1 proteins were expressed and purified in insect cells and their hormone induced interaction with *E. coli*-purified MBP-MpJAZ protein was tested in PD experiments. As shown in Figure 6B, dn-OPDA induces the interaction of the purified MpCOI1 and MpJAZ, similar to the effect of JA-Ile on Arabidopsis COI1-JAZs. In contrast, treatment with J4 substantially inhibited the dn-OPDA-dependent formation of the MpCOI1-MpJAZ complex (Figure 6B). We subsequently analysed the effect of J4 *in planta* by studying the OPDA-regulated transcriptional expression of several OPDA marker genes such as Mp*JAZ*, Mp*DIR*, Mp*PAT* and Mp*BHLH4* (Monte et al., 2018; Monte et al., 2019; Peñuelas et al., 2019). Indeed, exogenous pre-treatment with J4 partially prevents the transcriptional activation of these marker genes by OPDA (Figure 6C and Suppl. Figure S13).

**Figure 6.**
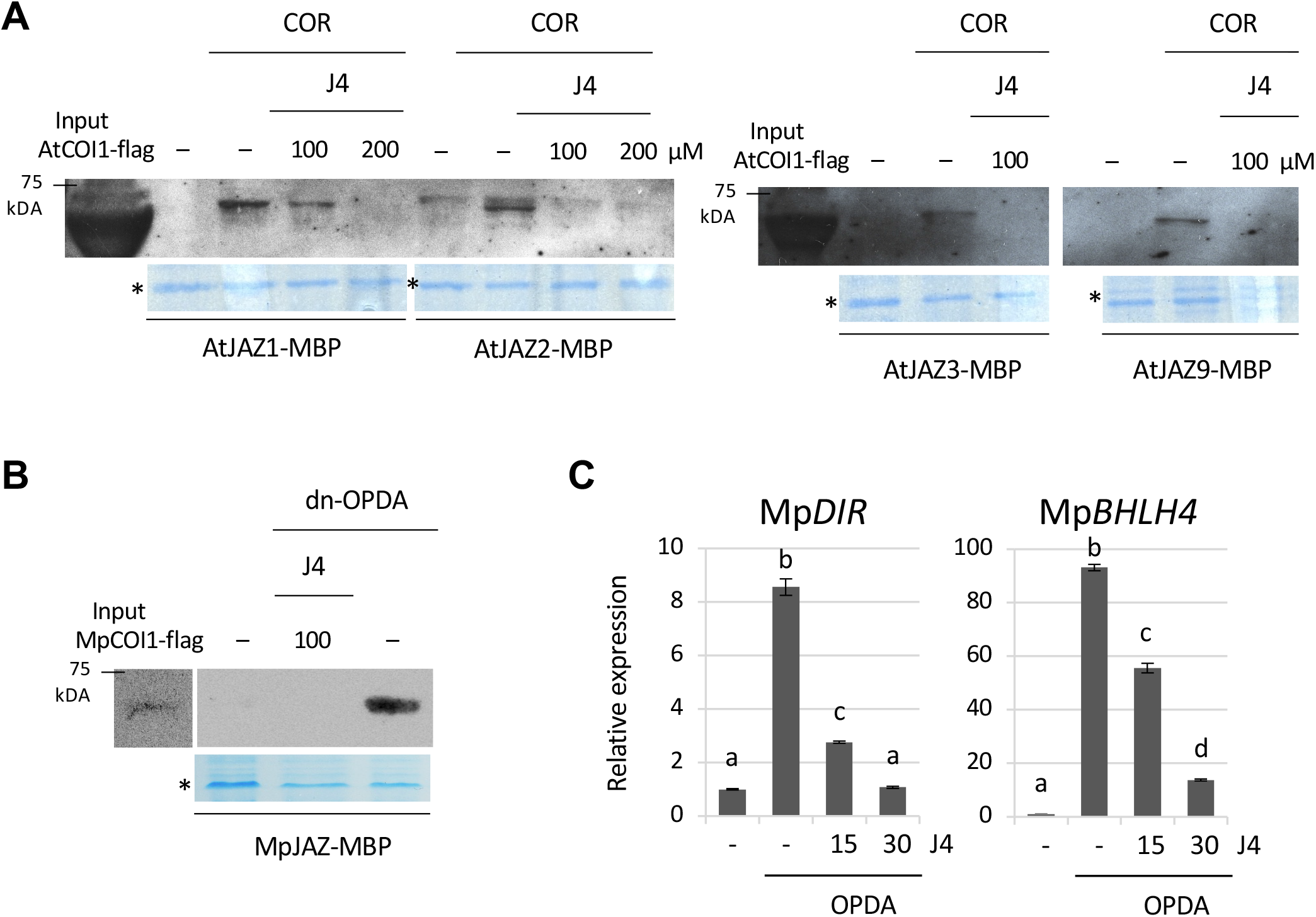
**J4 interferes with the hormone-dependent formation of several COI1- JAZ receptor complexes.** (A) Immunoblot analysis of AtCOI1-flag/AtJAZ-MBP interaction performed with anti- flag antibodies to COI1-flag proteins recovered from MBP-AtJAZ columns. ASK1- flag and COI1-flag were expressed in Sf9 insect cells, whereas AtJAZ were expressed in *E. coli*. The AtCOI1-flag/AtJAZ-MBP interaction is dependent on the presence of COR (0.5 μM); the compound J4 (at 100 and 200 μM) inhibits the hormone-dependent AtCOI1-flag/AtJAZ-MBP interaction. J4 was dissolved in DMSO; therefore, an equivalent volume of DMSO was used as negative control (labelled as -) in A-C. The lower panel shows Coomassie blue staining of the AtJAZ-MBP after cleavage with Factor Xa. Asterisks indicate the JAZ-MBP bands of the expected size in the Coomassie panel of each pull-down assay. (B) Immunoblot (antiflag antibody) of recovered MpCOI1-flag after pull-down reactions using recombinant MpJAZ-MBP protein alone (-) or with dinor-OPDA. MpASK1-flag and MpCOI1-flag were expressed in Sf9 insect cells, and MpJAZ was expressed in *E. coli*. The MpCOI1-flag/MpJAZ-MBP interaction depends on the presence of dn-OPDA (50 μM). The J4 compound (at 100 and 200 μM) inhibits the hormone-dependent MpCOI1-flag/MpJAZ-MBP interaction. Bottom, Coomassie blue staining of MpJAZ-MBP after cleavage with Factor Xa. (C) Gene expression analysis of Mp*DIR* (Dirigent-like protein) and Mp*BHLH4* (basic helix-loop-helix 14) in Tak-1 Marchantia plants in response to 10 µM OPDA for 1 hour; plants pre-treated with DMSO (-, untreated control), 15 or 30 µM J4 (+) for 1 hour. Mp*ACT* was used as housekeeping control gene. One-way ANOVA with post-hoc Tukey HSD Test (p< 0.01) analyses define the significant differences in gene expression. Each biological sample consisted of tissue pooled from 5-8 plants. Data show mean ± SD of four technical replicates.

Altogether, these results show that J4 directly interferes with the hormone-triggered establishment of the COI1-JAZ receptor complexes of vascular and non-vascular plants in absence of any other plant proteins. In addition, J4 has an *in vivo* inhibitory effect on the jasmonates-mediated responses in both vascular and non-vascular plants. Finally, the absence of putative regulators of COI1 or JAZ proteins in these assays (Figure 6A-B) indicate that J4 has a direct effect on the formation of the hormone-promoted COI1-JAZ complexes, likely competing with the ligands for the binding pocket.

## Discussion

To date, the role of JA regulating plant development and adaptation to biotic and abiotic stresses has been primarily analysed using classical genetic approaches relying on mutants impaired in the biosynthesis or response to JA (Browse 2009, Chini*, et al.* 2016). However, these loss-of-function genetic tools hold some intrinsic limitations such as functional redundancy, sterility and lethality (Xie*, et al.* 1998, Sanders *et al*. 2000, Browse 2009, Fernandez-Calvo*, et al.* 2011, Qi*, et al.* 2015, Chini *et al*. 2018b). Thus, the identification of molecules that specifically prevent JA-Ile- perception represent excellent tools to explore the role of JA in specific plant stages or tissues, avoiding these limitations. For example, specific auxin inhibitors were critical tools to understand the role of auxins in various stages of plant development and different plant species (Hayashi*, et al.* 2012, Leyser 2018). Using the auxin inhibitor naxillin researchers defined a previously unknown function for root cap in root branching by specifically stimulating the accumulation of active endogenous auxin only in the root cap (De Rybel *et al*. 2012).

Molecules antagonizing JA-perception would allow the manipulation of the JA pathway in a specific tissue or developmental stage in a reversible manner. Ideally, the antagonists identified in Arabidopsis should be easily transferred to other plant species and might clarify the role of JA-Ile in different plants. The two-step approach reported here strongly minimizes the identification of false hits, one of the most problematic constraints in chemical screens (Hicks and Robert 2014, Serrano *et al*. 2015). Thus, we identified only three molecules (among the more than 20,000 compounds screened) with robust JA-Ile antagonistic activity in both bioassays (Figure 1). Further biochemical and physiological analyses confirmed J4 as the only compound with robust JA-Ile antagonistic activity on several transcriptional and physiological JA-responses *in planta* at concentrations similar to that of JA (Figure 3 and Suppl. Figure S9).

The toxicity and secondary effects of current agrochemicals can seriously affect the environment and, potentially, animal and human health (Enserink *et al*. 2013, Lamberth*, et al.* 2013). Multiple, unspecific or pleiotropic effects are rather common among compounds identified in chemical screens (Fonseca *et al*. 2014b, Serrano*, et al.* 2015). Exceptionally, chemical screens also identified molecules with very specific targets and limited or none off-target effects. For example, Kynurenine, identified as inhibitor of the ethylene responses in root tissues (He*, et al.* 2011), is the first specific inhibitor of auxin biosynthesis binding to the substrate pocket of the TAA1/TAR aminotransferases family but no other related aminotransferases.

Here, we showed that J4 acts as a specific and dual inhibitor of the closely- related jasmonate and auxin signalling pathways by antagonizing the hormone-induced formation of the co-receptor complexes. Moreover, this inhibitory activity is evolutionarily conserved in land plants (at least between liverworts and eudicots) since J4 also inhibits the formation of the Marchantia MpCOI1-dn-OPDA-MpJAZ complex. Jasmonate and auxin pathways share strikingly similar receptors (COI1 and TIR1), which have a common evolutionary origin and share many conserved residues (Bowman *et al*. 2017). Thus, we propose that J4 can partially mimic dn-OPDA, JA-Ile and auxin to enter the COI1/TIR1 binding pocket, but cannot stablish the complex due to a lack of interaction with the JAZ/Aux-IAA co-receptors.

Therefore, J4 may be considered a specific antagonist of two related receptors that, however, do not affect other hormonal pathways or general proteasome-related mechanisms. Our results support the use of J4 as a commercially available inhibitor of JA- and auxin pathways in a particular tissue or developmental stage, useful in plant research and for agronomic purposes. J4 could also be useful in dissecting JA/auxin crosstalk. However, the concurrent inhibition of these two hormonal pathways may be undesirable in some circumstances. Therefore, additional structure activity relationships (SAR) analysis may be carried out to identify molecules structurally related J4 that potentially act as specific inhibitors of the JA- or auxin-pathway.

An alternative to chemical screen is the rational design of antagonist molecules specifically binding to the active pocket of key proteins. This approach, based on structural information of receptor-ligand complexes, is extensively exploited in medical research but just emerging in the agrochemical field (Lamberth*, et al.* 2013). For instance, the rational design of auxin analogues successfully obtained receptor antagonists useful in evolutionary distant plants species and that overcome the redundancy of auxin receptors (Hayashi *et al*. 2008, Hayashi*, et al.* 2012). Similarly, based on the crystal structure of the COI1-JAZ co-receptor, we designed a COR- derivative (COR-MO) with a very strong and specific activity preventing COI-JAZ interaction (Monte *et al*. 2014). Despite rational design being a very efficient and specific approach, it requires a deep knowledge of the structures of receptor-ligand complexes. In addition, the synthesis of rational-designed molecules is often very complex and expensive. A limited amount of compounds is therefore suitable only for specific, small-scale laboratory assays and not for agronomic use. Moreover, their specificity may prevent their use in different species. For instance, bryophytes such as *Marchantia polymorpha* do not synthesise JA-Ile, and the bioactive jasmonate and ligand of the Marchantia COI1-JAZ co-receptor is dn-OPDA (Monte *et al*. 2018). The Marchantia COI1-JAZ fails to bind JA-Ile or COR, and therefore, COR-MO cannot be used as JA-inhibitor in non-vascular plants (Monte*, et al.* 2018). In contrast, J4 is able to block all COI1-JAZ complexes tested (from Marchantia and Arabidopsis), suggesting that J4 is potentially active in all plants, from bryophytes to angiosperm. This broad activity of J4 underscores the high agronomic potential of this compound and its derivates because they can be active in all crops. Therefore, the “unbiased” chemical screen using commercially-available chemically diverse libraries is an informative approach.

Finally, the robustness of the reported screening system encourages to undertake additional screens using newly accessible “natural compound libraries” to identify natural molecules directly perturbing the COI1-JAZ co-receptor complex. Natural JA-Ile agonists could represent either novel activators of hormone biosynthesis or different active forms of the plant hormone, modulating specific COI1-JAZ complexes. Besides, the *in planta* optimized screen described here open the way to identify natural molecules affecting previously unknown post-transcriptional regulatory mechanisms of the key components of the JA signalling pathway, the JAZ repressors.

## Material & Methods

### Plant materials, growth conditions and transgenic plants

*Arabidopsis thaliana* Col-0 is the genetic background of wild type and transgenic lines used throughout the work. Seeds were surface sterilized and kept at 4°C in the dark for 48 h, then grown at 23°C with a 16-hour day cycle for 6 days, as previously described (Chini*, et al.* 2018b, Chico *et al*. 2020). *Nicotiana benthamiana* and tomato seeds were germinated and grown in the same conditions. Vertically grown seedlings were germinated in the same conditions. The transgenic line 35S:JAZ1-GUS was generated by (Thines*, et al.* 2007). The generation of transgenic plants expressing 35S:JAZ9-GUS were previously described (Monte *et al*. 2014). The *pJAZ9:GUS* and *pJAZ2:GUS* reporter lines were also previously described (Monte *et al*. 2014, Gimenez-Ibanez *et al*. 2017). The homozygous *pJAZ9:GUS* line was crossed with the loss-of- function *coi1-30* mutant and the double homozygous *pJAZ9:GUS coi1-30* line was isolated.

### Yeast-two-hybrid assays

The growth, handling and transformation of yeast were as previously described (Chini*, et al.* 2009b, Chini 2014). Briefly, the described plasmids were co-transformed into *Saccharomyces cerevisiae* AH109 cells following standard LiAC protocols to assess protein interactions. Successfully transformed colonies were identified on yeast synthetic dropout lacking Leu and Trp. 3 days after transformation, yeast colonies were grown in selective -WL liquid media for 6/7 hours and the cell density was adjusted to 3 x 10^7^ cells/ml (OD_600_ = 1). 4 μl of the cell suspensions were plated out on yeast synthetic dropout lacking Ade, His, Leu and Trp to test protein interaction (supplemented with coronatine and antagonist compounds as indicated). Plates were incubated at 28°C for 2 to 4 days. As positive control, the yeast suspensions were also plated on -AHWL media containing coronatine and the appropriate volume of DMSO.

### Chemical libraries and compounds

In the COI1-JAZ Y2H screen, we tested 22,680 compounds from three chemical libraries: the 2,320 molecules of MicroSource (from MDSI, http://www.msdiscovery.com/spectrum.html), the approximately 20,000 compounds of the 20 K DIVERSet™ (from ChemBridge, http://www.chembridge.com) and the 360 bioactive compounds described by (Drakakaki *et al*. 2011). In the JAZ1-GUS screen, compounds from the libraries MicroSource and 360 bioactive, and a subset of the 20 K DIVERSet™ were employed. All chemicals were stored at -20°C as 10 mg/ml stocks in 100% DMSO and supplemented to a final concentration of 50–100 μM based on the molecular mass of each compound.

### GUS staining and visualization

The visualization of GUS (β-glucuronidase) in the four JAZ1-GUS, JAZ9- GUS, *pJAZ2:GUS* and *pJAZ9:GUS* marker lines was carried out as previously described (Chini 2014, Chini *et al*. 2018a). JAZ1-GUS and JAZ9-GUS seeds were germinated in MS plates congaing 1% agar, kept in vertical for 7 days, transferred to 2 ml Johnson’s liquid media supplemented with the described chemicals and incubated in an orbital shaker at 100 rpm, as described in (Chini 2014). Seedlings of the *pJAZ9:GUS* and *pJAZ2:GUS* lines were germinated in liquid MS media in an orbital shaker at 100 rpm. Chemical treatments were carried out 6 days after germination.

After the described treatments, samples were placed in staining solution 50 mM phosphate buffer (pH 7), 0.1 % (v/v) Triton X-100 (Sigma), 1 mM X-Gluc (Glycosynth), 1 mM K-ferrocyanide (Sigma), 1 mM K-ferricyanide (Sigma) and incubated at 37°C overnight. After staining, the seedlings were washed in of 75% ethanol to eliminated chlorophyll. JAZ1-GUS seedlings were otherwise treated as described above and roots were collected. Protein extract was employed for fluorometric quantification of GUS activity using a Spectra Max M2 flouorometer (Molecular Devices) (Monte*, et al.* 2014).

### Degradation of RGA-GFP and EIN3-GFP

pRGA:GFP-RGA, 35S:EIN3-GFP and 35S:HA-IAA1 (Silverstone*, et al.* 2001, Guo and Ecker 2003) seeds were germinated in 2 ml liquid MS media in an orbital shaker at 100 rpm and the described compounds were added directly to the media 6 days after germination. After 2 hours, 20 plants of the same size were collected and proteins were extracted as previously described (Fonseca and Solano 2013). Samples were denatured, loaded on 9% SDS–PAGE gels, transferred to nitrocellulose membranes (Bio-Rad) and incubated with anti-GFP-HRP antibody (Milteny Biotec). Blots were developed using ECL (Pierce).

### Protein extracts and pull-down assays

MBP-JAZ fusion proteins were generated as previously described (Fonseca*, et al.* 2009). Pull-down experimental procedure is extensively described by (Fonseca and Solano 2013). Briefly, 35:COI1-flag seedlings were ground in liquid nitrogen and homogenized in extraction buffer. For PD experiments, 6 μg of resin-bound MBP fusion protein was added to 1 mg of total protein extract and incubated for 1 hour at 4 °C with rotation in presence of coronatine and the indicated compounds or DMSO control. After washing, samples were denatured, loaded on 8% SDS–PAGE gels, transferred to nitrocellulose membranes and incubated with anti-flag antibody (Sigma). To confirm equal protein loading, 7 µl of MBP-fused protein of each sample was run into SDS-PAGE gels and stained with Coomassie blue.

### Recombinant protein expression by baculovirus–insect cell expression systems

The expression of Arabidopsis and Marchantia ASK1 and COI1 in Sf9 insect cells was carried out as previously described (Sheard *et al*. 2010; Li *et al*. 2017; Takaoka *et al*. 2018). Briefly, pFast-HIS-flag-COI1 and pFastBac1-flag-ASK1 vectors were transformed into DH10Bac competent cell and further transposed into the bacmid. Blue/white selection were used to identify colonies containing the recombinant bacmid. The recombinant bacmid was transfected into Sf9 insect cells using Lipofectamine (Invitrogen) according to the manufacturers’ instructions. After the titter check, the isolated P1 recombinant baculovirus was further amplified to generate P2 and P3 stocks. P3 recombinant baculoviruses were added to a large cultured Sf9 cells at a volume ratio of 4.5:1000. 50 µl of insect cell culture supernatant for ASK1 and 50 µl for AtCOI1 or 50 µl for MpCOI1 were employed, together with 100 µl of PD buffer, for pull-down assays in presence of AtJAZ-MBP or MpJAZ proteins expressed in *E. coli* as reported above and extensively described by (Fonseca and Solano 2013).

### Root measurement, Anthocyanin and Chlorophyll quantification

Root growth inhibition assay was carried as previously described (Fonseca *et al*. 2014a). 3-day-old Arabidopsis seedlings germinated in vertical plates were transferred onto vertical Johnson medium in presence of JA with or without 5 μM J4 for 10 days and root length of 15 to 50 seedlings was measured. Roots were quantified using ImageJ software. Values represent mean and the error bars standard deviation.

The same growing conditions were employed to measure chlorophylls using 13- day-old Arabidopsis seedlings grown for10 days in presence 10 μM JA with or with 5 μM J4. *Nicotiana benthamiana* seeds were germinated and grown at 23°C in a 16-hour day cycle. *N. benthamiana* seedlings were germinated and grown in vertical MS plates for 7 days; seedlings were then transferred in 5 ml liquid MS media supplemented with 25 μM JA with or without 10 μM J4. Three days later, the aerial part of 5 to 7 seedlings was collected for each measurement and 4 to 6 independent replicates were analysed. Chlorophyll measurements were performed as previously described (Fonseca*, et al.* 2014a). Acetone 80% (V/V) was used for extraction and absorbance at 645 and 663 nm was measured in a spectrophotometer (Spectra Max M2 Molecular Devices). For carotenoids measurements, samples were extracted overnight in 95% ethanol and absorbance at 470, 645 and 663 nm was measured in a spectrophotometer. Values represent mean and the error bars correspond to standard deviation. The experiment was repeated three with similar results.

Anthocyanin quantification was carried as previously described (Fonseca*, et al.* 2014a). Briefly, seedlings grown in vertical Johnson plates for 5 days and transferred to liquid Johnson media supplemented with 10 μM JA with or without 2.5 μM J4. Two days later, the aerial parts of 10 to 15 seedlings were pooled for each replicate and anthocyanin quantification was performed using a spectrophotometer. Four independent replicates (seedling pools) were measured for each sample. Values represent mean and the error bars standard deviation. The experiment was repeated at least three times with similar results.

### Quantitative RT-PCR

Quantitative RT-PCR was performed using biological samples of tissue pooled from 10–15 Arabidopsis seedlings. RNA was extracted and purified using a Plant Total RNA isolation kit (Favorgen), including on-column DNase digestion to remove genomic DNA contamination. cDNA was synthesized from 1 μg total RNA with a high-capacity cDNA reverse transcription kit (Applied Biosystems). Gene amplification was caried out using 4 μl from a 1:10 cDNA dilution, 4 μl of EvaGreen® qPCR Mix Plus (Solis BioDyne) and gene-specific primers (Supplemental Table S2). Quantitative PCR was performed in 96-well optical plates in a HT 7900 Real Time PCR system (Applied Biosystems) using standard thermocycler conditions. Relative expression values given as the means of three or four technical replicates (from 10–15 Arabidopsis seedlings) relative to the mock wild-type control using ACT8 as the housekeeping gene. Data were analysed using One-way ANOVA with post-hoc Tukey HSD Test (p< 0.01).

### Accession Numbers

Sequence data from this article can be found in the GenBank/EMBL data libraries under accession numbers_. *AtJAZ1*/*AT1G19180* (gene ID 838501), *AtJAZ3*/*AT3G17860* (821055), *AtCOI1*/*AT2G39940* (818581), *MpJAZ*/*Mapoly0097s0021/Mp6g06230* (P535) and *MpCOI1*/*Mapoly0025s0025/ Mp2g26590* (P322).

## Supplemental Data

**Supplemental Figure S1. Inhibition of COR-induced COI1-JAZ interactions in yeast two- hybrid assays.**

**Supplemental Figure S2. Molecules antagonizing the JA-mediated degradation of JAZ1-GUS protein *in planta*.**

**Supplemental Figure S3. Confirmation of the antagonistic effect of the 12 identified molecules on COI1-JAZ interaction in Y2H and JAZ1 degradation *in planta*.**

**Supplemental Figure S4. Minimal concentrations of antagonist molecules required to prevent COR-mediated COI1/JAZ interactions in Y2H.**

**Supplemental Figure S5. Quantification of the inhibition of JAZ1-GUS degradation by antagonist molecules *in vivo*.**

**Supplemental Figure S6. Inhibition of COI1-JAZ interactions in Y2H and of JAZ degradation *in planta* by derivate molecules of J4 and Y20.**

**Supplemental Figure S7. Inhibition of *JAZ2 and JAZ9* expression of derivate molecules of J4 and Y20.**

**Supplemental Figure S8. Specific expression of *JAZ9* in trichomes requires COI1 and it is inhibited by J4 treatment.**

**Supplemental Figure S9. J4 partially inhibits JA-mediated transcriptional activation in several angiosperm plants.**

**Supplemental Figure S10. Y11 and Y20 do not prevent JA-mediated responses *in planta*.**

**Supplemental Figure S11. Half maximal inhibitory concentration (IC50) of J4 in JA-promoted JAZ1-GUS degradation.**

**Supplemental Figure S12. Antagonistic effect of the compound J4 on JA-induced responses in tobacco plants.**

**Supplemental Figure S13. J4 partially inhibits OPDA-mediated transcriptional activation.**

**Supplemental Table S1. List of all identified molecules including their basic information.**

**Supplemental Table S2. List of primers sequences used for qPCR analyses.**

## Acknowledgments

We thank John Browse, Salome Prat, Joe Ecker and Judy Callis for providing 35S:JAZ1-GUS, pRGA:GFP-RGA, 35S:EIN3-GFP and 35S:HA-IAA1 seeds respectively. We also thank Daoxin Xie for providing the pFast-HIS-FLAG-COI1 and pFastBac1-FLAG-ASK1 vectors for expression in insect cells. We are grateful to Monica Diez Diaz for the selection of JAZ9-GUS transgenic line, Abel Rosado, David Carter and Michelle Brown for the technical help setting-up the chemical screen and Guillermo Gimenez Aleman for the structure images using ChemDraw. This work was financed by grants to R.S. and A.C. (BIO2013-44407-R and BIO2016-77216-R from MINECO/FEDER, and PID2019-107012RB-I00 from the Spanish Ministry of Science and Innovation AEI/FEDER) and the DE-FG02-11ER15295 grant to N.V.R. and G.R.H. from the US Department of Energy. A.C. was supported by the Ramon y Cajal (RYC-2010-05680) and HSPO Fellowships. I.M. was supported by a master fellowship from Fundación Ramón Areces/UAM and a predoctoral fellowship from the Ministerio de Educación (AP2010-1410). M.B. was supported by a JAE-DOC fellowship (CSIC).

## Supplementary Material

Supplemental Figure S1. **Inhibition of COR-induced COI1-JAZ interactions in yeast two- hybrid assays.**

Yeast cells co-transformed with pGAD-JAZ9 or pGAD-JAZ3 (preys) and pGBK- COI1, pGBK-NINJA or pGBK-JAZ9 (baits) were grown on yeast synthetic drop-out lacking Leu and Trp (-LW), as control or on selective media lacking Ade, His, Leu and Trp (-AHLW), to test protein interactions. COI1 interaction with JAZ9 (A) and JAZ3

(B) is detected only in presence 5 and 20 μM COR respectively. This figure shows the Y2H interaction in presence of COR and the indicated antagonist molecules (Y10 at 100 μM; Y17 at 25 μM; Y18 at 20 μM; Y20 at 15 μM; Y11 at 300 nM). Compounds were dissolved in DMSO, therefore as negative control an equivalent volume of DMSO was used (labelled as -). As control, we tested that the antagonist molecules did not interfere with the interaction between JAZ9 and NINJA or JAZ9 dimerization.

Supplemental Figure S2. **Molecules antagonizing the JA-mediated degradation of JAZ1-GUS protein *in planta*.**

Roots of 7-day-old JAZ1-GUS seedlings concurrently treated with 5 μM JA and 100 μM of the indicated molecules (J1, J2, J3, J4, J9, J10, J11) for 1 hour. JA treatment triggers the degradation of JAZ1-GUS, whereas the addition of the identified molecules could prevent JAZ1 destabilization. Compounds were prepared in DMSO; as negative control contained an equivalent volume of DMSO (defined as -).

The original chemical screen was carried out using COR whereas the JA was employed in the secondary confirmation assays to discard unspecific effects of COR or an effect of the compound on JA conversion into JA-Ile.

Supplemental Figure S3. **Confirmation of the antagonistic effect of the 12 identified molecules on COI1-JAZ interaction in Y2H and JAZ1 degradation *in planta*.**

(A-B) Yeast cells co-transformed with pGAD-JAZ9 or pGAD-JAZ3 (preys) and pGBK-COI1, pGBK-NINJA or pGBK-JAZ9 (baits) were selected and subsequently grown on yeast synthetic drop-out lacking Leu and Trp (-LW), as a transformation control or on selective media lacking Ade, His, Leu and Trp (-AHLW), to test protein interactions. COI1 interaction with JAZ9 (A) and JAZ3 (B) is induced only in presence 5 and 20 μM COR respectively. This figure shows the Y2H interaction in presence of COR and the indicated antagonist molecules (J1 and J9 at 100 μM; J10 at 50 μM; J4 at 15 μM; J2 at 10 μM; J3 at 10 nM, J11 at 5 nM). As control, we tested that the antagonist molecules did not interfere with the interaction between JAZ9 and NINJA or JAZ9 dimerization. Compounds were dissolved in DMSO, therefore as negative control an equivalent volume of DMSO was used (labelled as -) in A-C.

(C) Roots of 7-day-old JAZ1-GUS seedlings concurrently treated with 2 μM JA and 100 μM of the indicated molecules (Y10, Y11, Y17, Y18, Y20) for 1 hour. JA treatment triggers the degradation of JAZ1-GUS, whereas the addition of most antagonist molecules could prevent JAZ1 destabilization.

Supplemental Figure S4. **Minimal concentrations of antagonist molecules required to prevent COR-mediated COI1/JAZ interactions in Y2H.**

Yeast cells co-transformed with pGAD-JAZ9 (prey) and pGBK-COI1, pGBK-NINJA or pGBK-JAZ9 (baits) were selected and subsequently grown on yeast synthetic drop- out lacking Leu and Trp (-LW), as a transformation control or on selective media lacking Ade, His, Leu and Trp (-AHLW), to test protein interactions. COI1-JAZ9 interaction is detected only in presence 5 μM coronatine. A range of concentration of antagonistic molecules was used to define the minimal concentration required to inhibit COI1-JAZ interaction: 25 μM J4 (A), 300 nM Y11 (B) and 5 μM Y20 (C). Compounds were dissolved in DMSO; negative controls carry an equivalent volume of DMSO was used (defined as -) in A-C. As control, we tested that these molecules did not interfere with the interaction between JAZ9 and NINJA or JAZ9 dimerization.

Supplemental Figure S5. **Quantification of the inhibition of JAZ1-GUS degradation by antagonist molecules *in vivo*.**

Twenty to thirty 7-day-old seedlings were incubated in medium containing 50 nM COR with or without indicated compounds for 1 hour. Fluorometric GUS quantification of roots of Arabidopsis JAZ1-GUS line is shown. Relative GUS activity is shown normalizing 100% value to mock treatment (absence of COR). Compounds were dissolved in DMSO, therefore as negative control an equivalent volume of DMSO was used (labelled as -). Columns represent mean of 6 readings and error bars are standard deviations. One-way ANOVA with post-hoc Tukey HSD Test (p< 0.01) analyses define the significant differences in JAZ1-GUS degradation.

Supplemental Figure S6. **Inhibition of COI1-JAZ interactions in Y2H and of JAZ degradation *in planta* by derivate molecules of J4 and Y20.**

(A) Chemical structures of the described J4 and Y20 molecules. Oxygen atoms are highlighted in red, nitrogen in blue, sulphur in yellow and fluorine in purple.

(B) COR induces the interaction between the COI1 receptor and JAZ co-receptors in Y2H assay, whereas most Y20 derivate molecules prevent the COR-dependent COI1- JAZ interaction. Yeast cells co-transformed with pGAD-JAZ9 or pGAD-JAZ3 (preys) and pGBK-COI1, pGBK-NINJA or pGBK-JAZ9 (baits) were grown on yeast synthetic drop-out lacking Leu and Trp (-LW), as control or on selective media lacking Ade, His, Leu and Trp (-AHLW), to test protein interactions. COI1 interaction with JAZ9 and JAZ3 is detected only in presence 5 and 20 μM coronatine respectively. Y20 derivate molecules were employed at 25 μM. As control, we tested that these molecules did not interfere with the interaction between JAZ9 and NINJA or JAZ9 dimerization. Compounds were prepared in DMSO; as negative control an equivalent volume of DMSO was used (labelled as -).

(C) The figure show roots of 6-day-old JAZ1-GUS and JAZ9-GUS seedlings concurrently treated with 2 μM jasmonic acid and 100 μM of the indicated molecules. JA induces the degradation of JAZ1 and JAZ9, whereas most J4 and Y20 derivate molecules could prevent JAZ degradation.

Supplemental Figure S7. **Inhibition of *JAZ2 and JAZ9* expression of derivate molecules of J4 and Y20.**

(A) Roots of seedlings of the *pJAZ2:GUS* marker line concurrently treated for 75 minutes with 5 μM JA and the indicated molecules (J4, Y20, Y20-L1 and Y20-L2 at 10 μM; J4-L1, J4-L2 and J4-L3 at 25 μM; Y20-L3 at 100 μM). JA triggers the expression of *JAZ2*, whereas most molecules could prevent the *JAZ* transcriptional activation. Compounds were dissolved in DMSO, therefore as negative control an equivalent volume of DMSO was used (labelled as -).

(B) Seedlings of the *pJAZ9:GUS* line were concurrently wounded and treated for 3 hours the indicated molecules (100 μM). Mechanical wounding induces the accumulation of endogenous JA-Ile and the expression of *JAZ9*, whereas the addition of most molecules could inhibit the transcriptional activation of *JAZ9*.

Supplemental Figure S8. **Specific expression of *JAZ9* in trichomes requires COI1 and it is inhibited by J4 treatment.**

(A-B) Leaves of 10-day-old *pJAZ9:GUS* seedlings in WT (A) and *coi1-30* (B) background are shown. Expression of *JAZ9* in trichomes requires *COI1*.

(C) *pJAZ9:GUS* seedlings were treated over-night with the indicated compounds at 100 μM. Only the J4 compound can inhibit the trichome-specific *JAZ9* expression. Compounds were prepared in DMSO, so an equivalent volume of DMSO was used as negative control (labelled as -).

Supplemental Figure S9. **J4 partially inhibits JA-mediated transcriptional activation in Arabidopsis.**

Gene expression analysis of *JAZ10*, *OPR3* and *TAT3* in wild-type (Col-0) plants in response to 12.5 µM JA for 45 minutes; plants pre-treated with DMSO (-; untreated control) or J4 (+) for 1 hour. *ACT8* was used as housekeeping control gene. One-way ANOVA with post-hoc Tukey HSD Test (p< 0.01) analyses define the significant differences gene expression. Each biological sample consisted of tissue pooled from 10-15 plants. Data show mean ± SD of three to four technical replicates. (A),

Supplemental Figure S10. **Y11 and Y20 do not prevent JA-mediated responses *in planta*.**

(A) 13-day-old WT seedlings germinated in MS media for 3 days and then grown for 10 days in vertical plate in presence of 10 μM JA and 2.5 μM Y11 or Y20. Compounds were dissolved in DMSO, therefore an equivalent volume of DMSO was used as negative control (labelled as -) in. A-C. Lines represent 1 cm.

(B) Root growth inhibition by 15 μM JA of 13-day-old WT seedlings grown for 10 days in vertical plate in presence or absence of 2.5 μM Y11 or Y20. Results are expressed as mean +/- SD of 25-30 plants. One-way ANOVA with post-hoc Tukey HSD Test (p< 0.01) analyses define the significant differences.

(C) Anthocyanin accumulation, shown as Absorbance (530) per gram of plant fresh weigh (FW), in 8-day-old WT seedlings grown for 2 days in presence of 50 μM JA with or without 2.5 μM Y11 or Y20. Results are expressed as mean +/- SD. One-way ANOVA with post-hoc Tukey HSD Test (p< 0.01) analyses define the significant differences.

Supplemental Figure S11. **Half maximal inhibitory concentration (IC50) of J4 in JA-promoted JAZ1-GUS degradation.**

(A) Quantification of GUS activity (arbitrary unit U per μg protein per h) in roots of 7- d-old JAZ1-GUS plants. Seedlings (N =20 to 30) were pretreated with the indicated concentrations of J4 for 1 h and then treated with 15 μM JA for 1 hour. Results shown are the mean ± s.d. of seven replicates.

(B) Relative quantification of JAZ1-GUS. Untreated control was set as 100% of GUS activity in roots of 7-d-old JAZ1-GUS plants. Seedlings (N =20 to 30) were pretreated for 1 hour with the indicated concentrations of J4 and then with 15 μM JA for an additional hour. Results shown are the mean ± s.d. of seven replicates. The plain line connects the experimental values, whereas the dotted line is the trendline.

Supplemental Figure S12. **Antagonistic effect of the compound J4 on JA-induced responses in *Nicotiana benthamiana* plants.**

Chlorophyll a and b (A) and carotenoids (B) content of 10-day-old *N. benthamiana* seedlings (N = 7 plants) germinated in vertical plates and then grown for 3 days in presence of 25 μM JA with or without 10 μM J4. The experiments were repeated three times with similar results.

A and B, Bars represent the average value and error bars the standard deviation. One-way ANOVA with post-hoc Tukey HSD Test (p< 0.01) analyses define the significant differences.

Supplemental Figure S13. **J4 partially inhibits OPDA-mediated transcriptional activation.**

Gene expression analysis of Mp*PAT* and Mp*JAZ* in Tak-1 Marchantia plants in response to 10 µM OPDA for 1 hour; plants pre-treated with DMSO (-; untreated control), 15 or 30 µM J4 (+) for 1 hour. Mp*ACT* was used as housekeeping control gene. One-way ANOVA with post-hoc Tukey HSD Test (p< 0.01) analyses define the significant differences in gene expression. Each biological sample consisted of tissue pooled from 5-8 plants. Data show mean ± SD of four technical replicates.

**Supplemental Table S1.** List of all identified molecules including their basic information.

**Supplemental Table S2.** List of primers sequences used for RT-qPCR analyses.

