## Supplemental Material for "A small molecule antagonizes jasmonic acid perception and auxin responses in vascular and non-vascular plants"

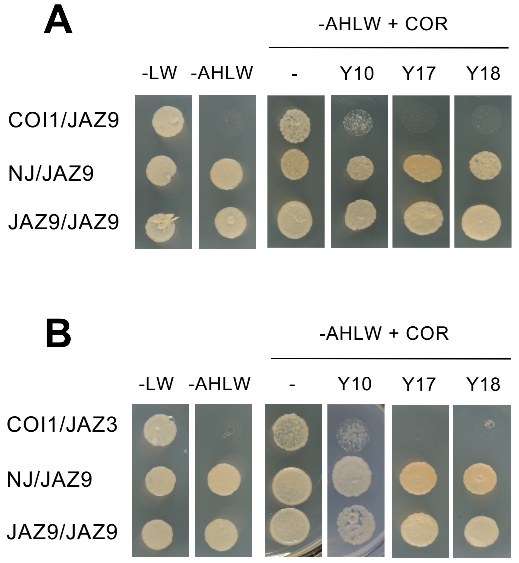


Supplemental Figure S1. **Inhibition of COR-induced COI1-JAZ interactions in yeast two- hybrid assays.**

Yeast cells co-transformed with pGAD-JAZ9 or pGAD-JAZ3 (preys) and pGBK-COI1, pGBK-NINJA or pGBK-JAZ9 (baits) were grown on yeast synthetic drop-out lacking Leu and Trp (-LW), as control or on selective media lacking Ade, His, Leu and Trp (-AHLW), to test protein interactions. COI1 interaction with JAZ9 (A) and JAZ3 (B) is detected only in presence 5 and 20 µM COR respectively. This figure shows the Y2H interaction in presence of COR and the indicated antagonist molecules (Y10 at 100 µM; Y17 at 25 µM; Y18 at 20 µM). Compounds were dissolved in DMSO, therefore as negative control an equivalent volume of DMSO was used (labelled as -). As control, we tested that the antagonist molecules did not interfere with the interaction between JAZ9 and NINJA or JAZ9 dimerization.

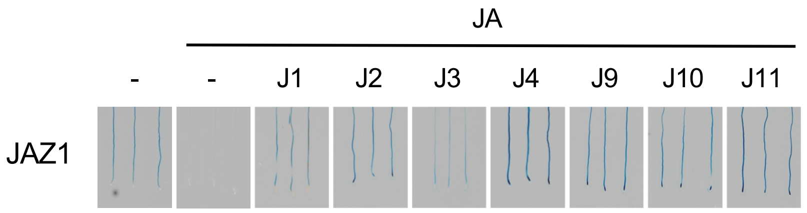


Supplemental Figure S2. **Molecules antagonizing the JA-mediated degradation of JAZ1-GUS protein *in planta***.

Roots of 7-day-old JAZ1-GUS seedlings concurrently treated with 5 µM JA and 100 µM of the indicated molecules for 1 hour. JA treatment triggers the degradation of JAZ1-GUS, whereas the addition of the identified molecules could prevent JAZ1 destabilization. Compounds were prepared in DMSO; as negative control contained an equivalent volume of DMSO (defined as -).

The original chemical screen was carried out using COR whereas the JA was employed in the secondary confirmation assays to discard unspecific effects of COR or an effect of the compound on JA conversion into JA-Ile.

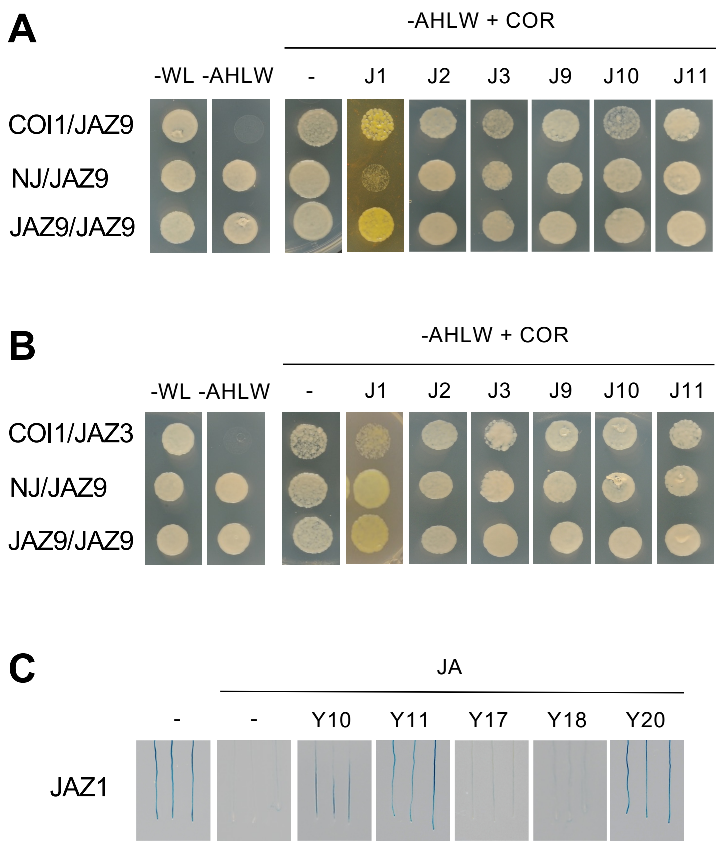


Supplemental Figure S3. **Confirmation of the antagonistic effect of the 12 identified molecules in COI1-JAZ interaction on Y2H and JAZ1 degradation *in planta***.

(A-B) Yeast cells co-transformed with pGAD-JAZ9 or pGAD-JAZ3 (preys) and pGBK-COI1, pGBK-NINJA or pGBK-JAZ9 (baits) were selected and subsequently grown on yeast synthetic drop-out lacking Leu and Trp (-LW), as a transformation control or on selective media lacking Ade, His, Leu and Trp (-AHLW), to test protein interactions. COI1 interaction with JAZ9 (A) and JAZ3 (B) is induced only in presence 5 and 20 µM COR respectively. This figure shows the Y2H interaction in presence of COR and the indicated antagonist molecules (J1 and J9 at 100 µM; J10 at 50 µM; J2 at 10 µM; J3 at 10 nM, J11 at 5 nM). As control, we tested that the antagonist molecules did not interfere with the interaction between JAZ9 and NINJA or JAZ9 dimerization. Compounds were dissolved in DMSO, therefore as negative control an equivalent volume of DMSO was used (labelled as -) in A-C.

(C) Roots of 7-day-old JAZ1-GUS seedlings concurrently treated with 2 µM JA and 100 µM of the indicated molecules for 1 hour. JA treatment triggers the degradation of JAZ1-GUS, whereas the addition of most antagonist molecules could prevent JAZ1 destabilization.

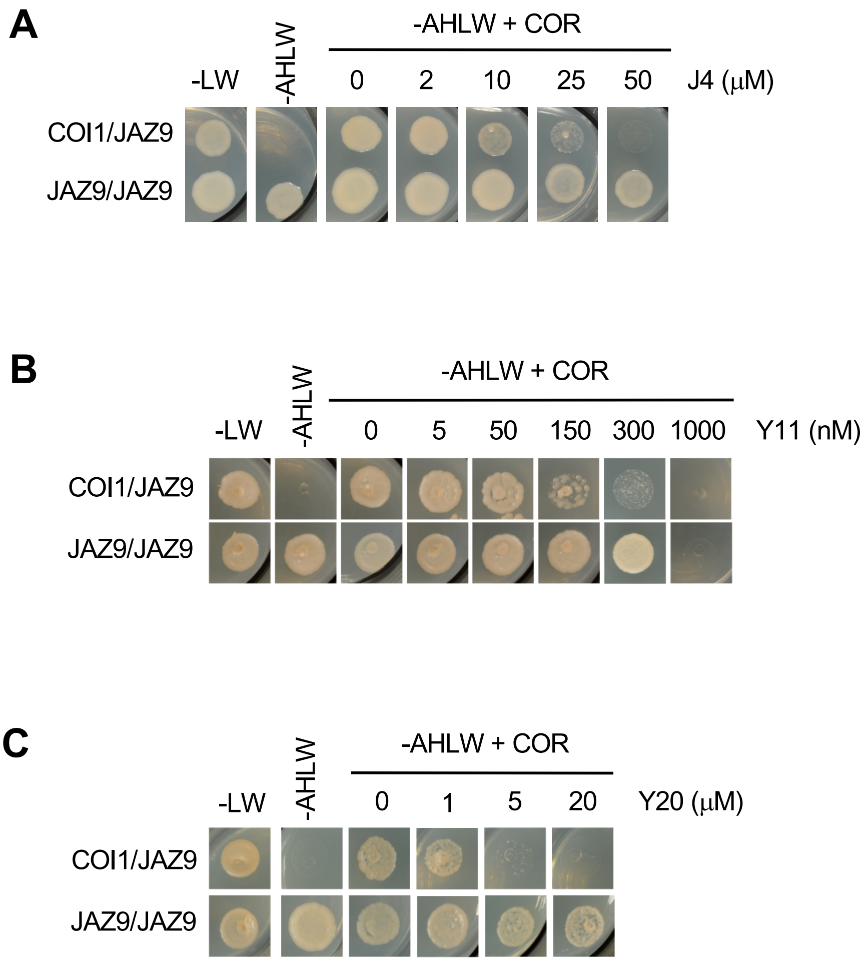


Supplemental Figure S4. **Minimal concentrations of antagonist molecules required to prevent COR-mediated COI1/JAZ interactions in Y2H**.

Yeast cells co-transformed with pGAD-JAZ9 (prey) and pGBK-COI1, pGBK-NINJA or pGBK-JAZ9 (baits) were selected and subsequently grown on yeast synthetic drop-out lacking Leu and Trp (-LW), as a transformation control or on selective media lacking Ade, His, Leu and Trp (-AHLW), to test protein interactions. COI1-JAZ9 interaction is detected only in presence 5 µM coronatine. A range of concentration of antagonistic molecules was used to define the minimal concentration required to inhibit COI1-JAZ interaction: 25 µM J4 (A), 300 nM Y11 (B) and 5 µM Y20 (C). Compounds were dissolved in DMSO; negative controls carry an equivalent volume of DMSO was used (defined as -) in A-C. As control, we tested that these molecules did not interfere with the interaction between JAZ9 and NINJA or JAZ9 dimerization.

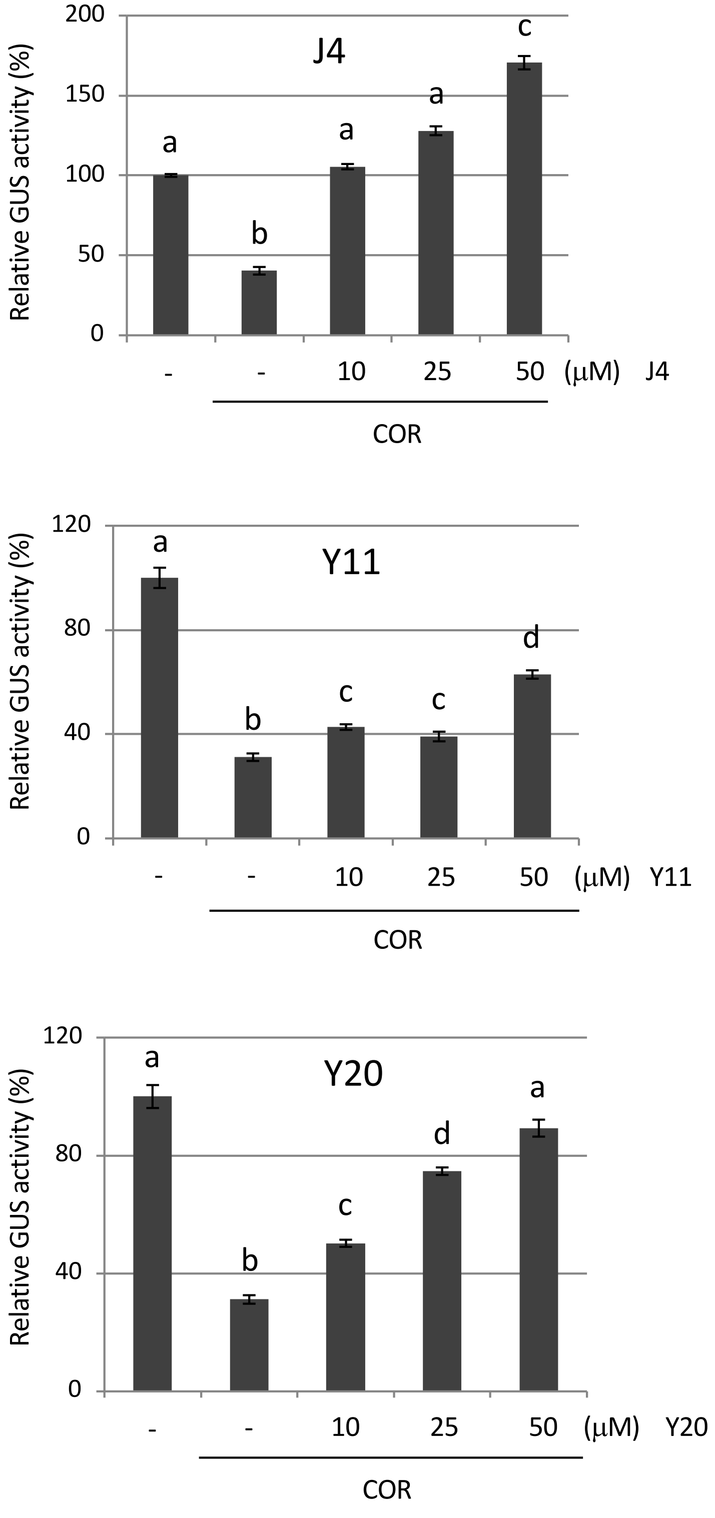


Supplemental Figure S5. **Quantification of the inhibition of JAZ1-GUS degradation by antagonist molecules *in vivo*.**

Twenty to thirty 7-day-old seedlings were incubated in medium containing 50 nM COR with or without indicated compounds for 1 hour. Fluorometric GUS quantification of roots of Arabidopsis JAZ1-GUS line is shown. Relative GUS activity is shown normalizing 100% value to mock treatment (absence of COR). Compounds were dissolved in DMSO, therefore as negative control an equivalent volume of DMSO was used (labelled as -). Columns represent mean of 6 readings and error bars are standard deviations. One-way ANOVA with post-hoc Tukey HSD Test ($p < 0.01$) analyses define the significant differences in JAZ1-GUS degradation.

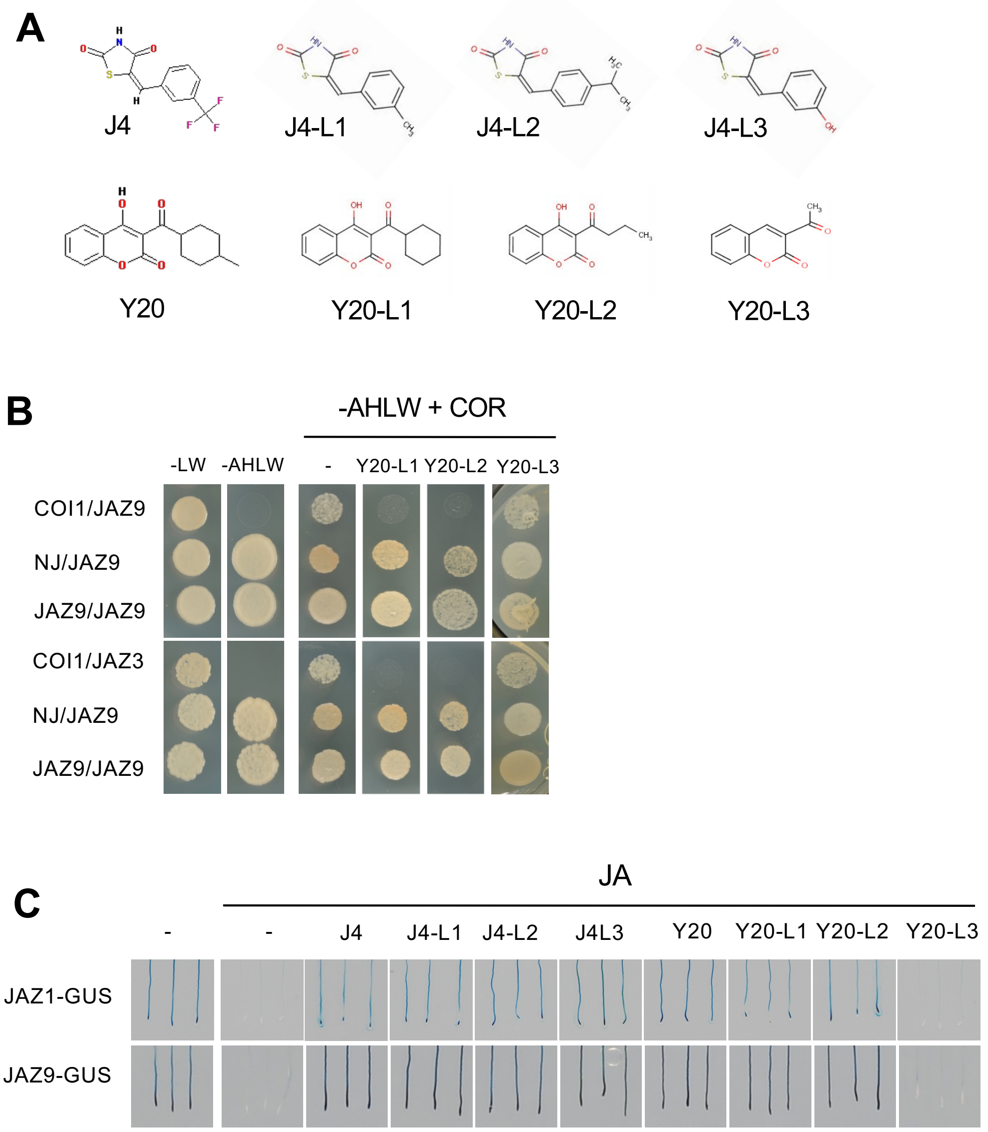


Supplemental Figure S6. **Inhibition of COI1-JAZ interactions in Y2H and of JAZ degradation *in planta* by derivate molecules of J4 and Y20**.

(A) Chemical structure of the described J4 and Y20 molecules.

(B) COR induces the interaction between the COI1 receptor and JAZ co-receptors in Y2H assay, whereas most Y20 derivate molecules prevent the COR-dependent COI1-JAZ interaction. Yeast cells co-transformed with pGAD-JAZ9 or pGAD-JAZ3 (preys) and pGBK-COI1, pGBK-NINJA or pGBK-JAZ9 (baits) were grown on yeast synthetic drop-out lacking Leu and Trp (-LW), as control or on selective media lacking Ade, His, Leu and Trp (-AHLW), to test protein interactions. COI1 interaction with JAZ9 and JAZ3 is detected only in presence 5 and 20 µM coronatine respectively. Y20 derivate molecules were employed at 25 µM. As control, we tested that these molecules did not interfere with the interaction between JAZ9 and NINJA or JAZ9 dimerization. Compounds were prepared in DMSO; as negative control an equivalent volume of DMSO was used (labelled as -).

(C) The figure show roots of 6-day-old JAZ1-GUS and JAZ9-GUS seedlings concurrently treated with 2 µM jasmonic acid and 100 µM of the indicated molecules. JA induces the degradation of JAZ1 and JAZ9, whereas most J4 and Y20 derivate molecules could prevent JAZ degradation. The J4 treatment of JAZ9-GUS is the same experiment shown in Figure 1C.

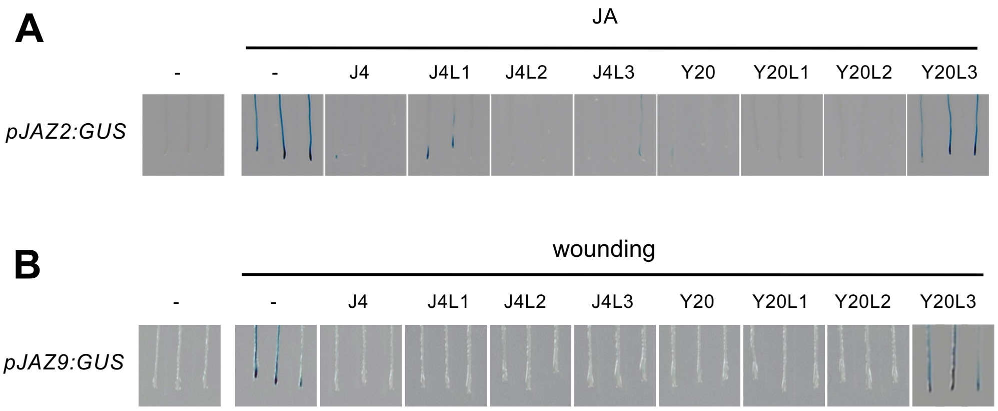


Supplemental Figure S7. **Inhibition of *JAZ2* and *JAZ9* expression of derivate molecules of J4 and Y20**.

(A) Roots of seedlings of the *pJAZ2:GUS* marker line concurrently treated for 75 minutes with 5 µM JA and the indicated molecules (J4, Y20, Y20-L1 and Y20-L2 at 10 µM; J4-L1, J4-L2 and J4-L3 at 25 µM; Y20-L3 at 100 µM). JA triggers the expression of *JAZ2*, whereas most molecules could prevent the *JAZ* transcriptional activation. . Compounds were dissolved in DMSO, therefore as negative control an equivalent volume of DMSO was used (labelled as -).

(B) Seedlings of the *pJAZ9:GUS* line were concurrently wounded and treated for 3 hours the indicated molecules (100uM). Mechanical wounding induces the accumulation of endogenous JA-Ile and the expression of *JAZ9*, whereas the addition of most molecules could inhibit the transcriptional activation of *JAZ9*.

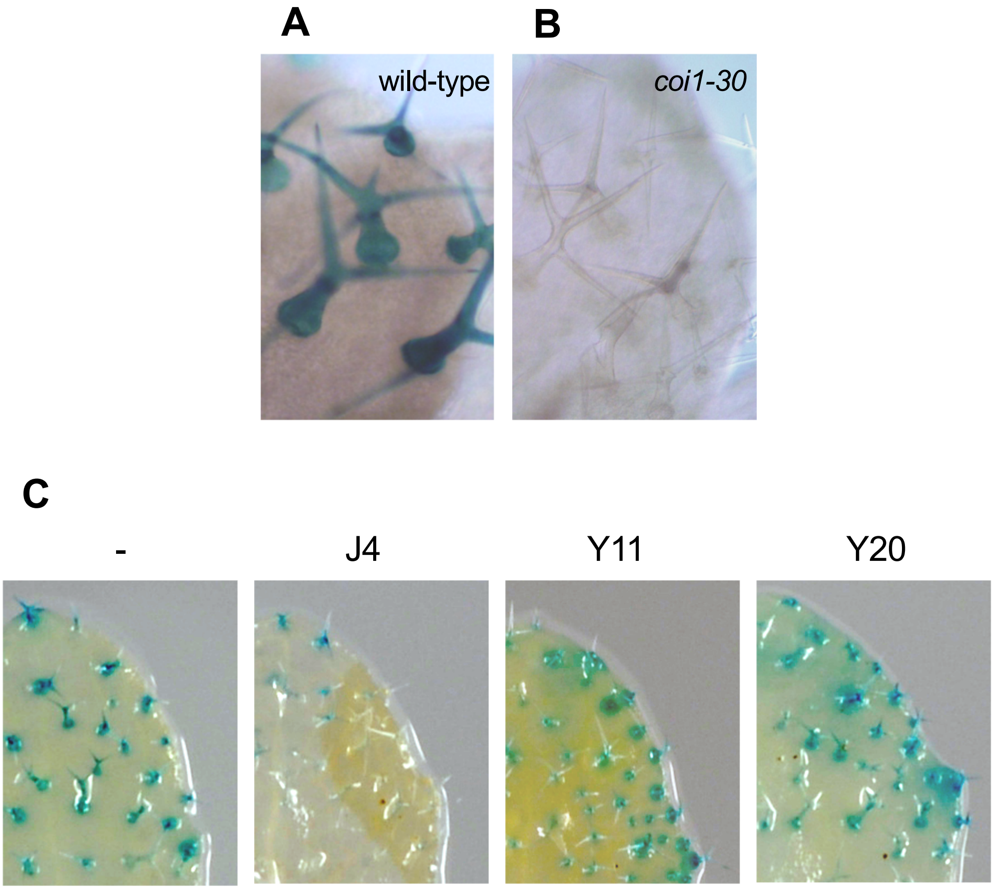


Supplemental Figure S8. **Specific expression of *JAZ9* in trichomes requires COI1 and it is inhibited by J4 treatment**.

(A-B) Leaves of 10-day-old *pJAZ9:GUS* seedlings in WT (A) and *coi1-30* (B) background are shown. Expression of *JAZ9* in trichomes requires *COI1*.

(C) *pJAZ9:GUS* seedlings were treated over-night with the indicated compounds at 100 μM. Only the J4 compound can inhibit the trichome-specific *JAZ9* expression. Compounds were prepared in DMSO, so an equivalent volume of DMSO was used as negative control (labelled as -).

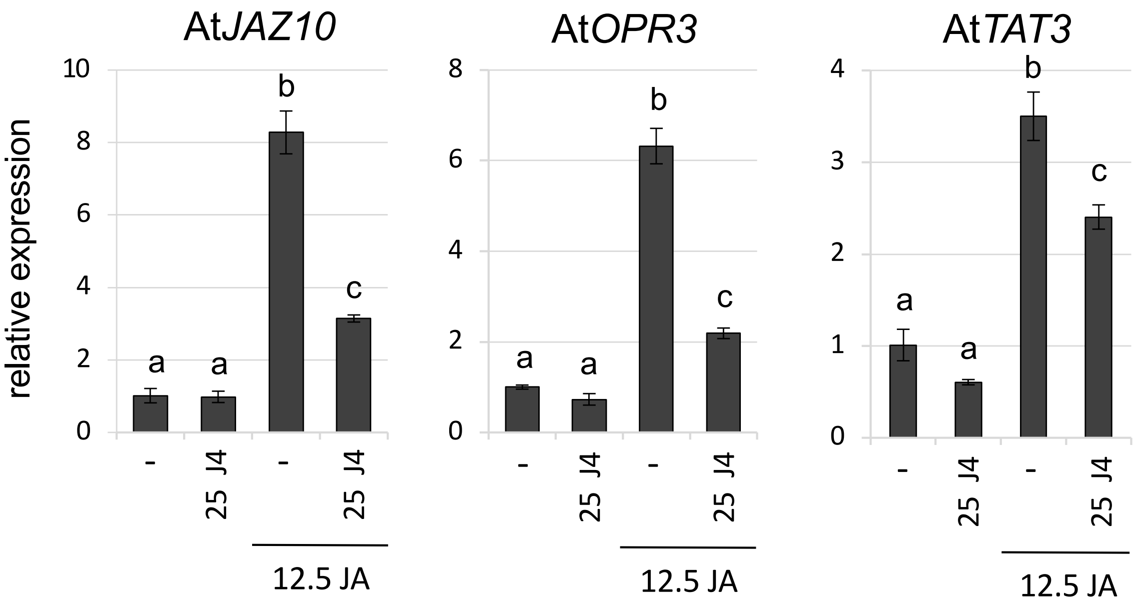


Supplemental Figure S9. **J4 partially inhibits JA-mediated transcriptional activation in Arabidopsis.**

Gene expression analysis of *JAZ10*, *OPR3* and *TAT3* in wild-type (Col-0) plants in response to 12.5 µM JA for 45 minutes; plants pre-treated with DMSO (-; untreated control) or J4 (+) for 1 hour. *ACT8* was used as housekeeping control gene. One-way ANOVA with post-hoc Tukey HSD Test ($p < 0.01$) analyses define the significant differences gene expression. Each biological sample consisted of tissue pooled from 10-15 plants. Data show mean ± SD of three to four technical replicates.

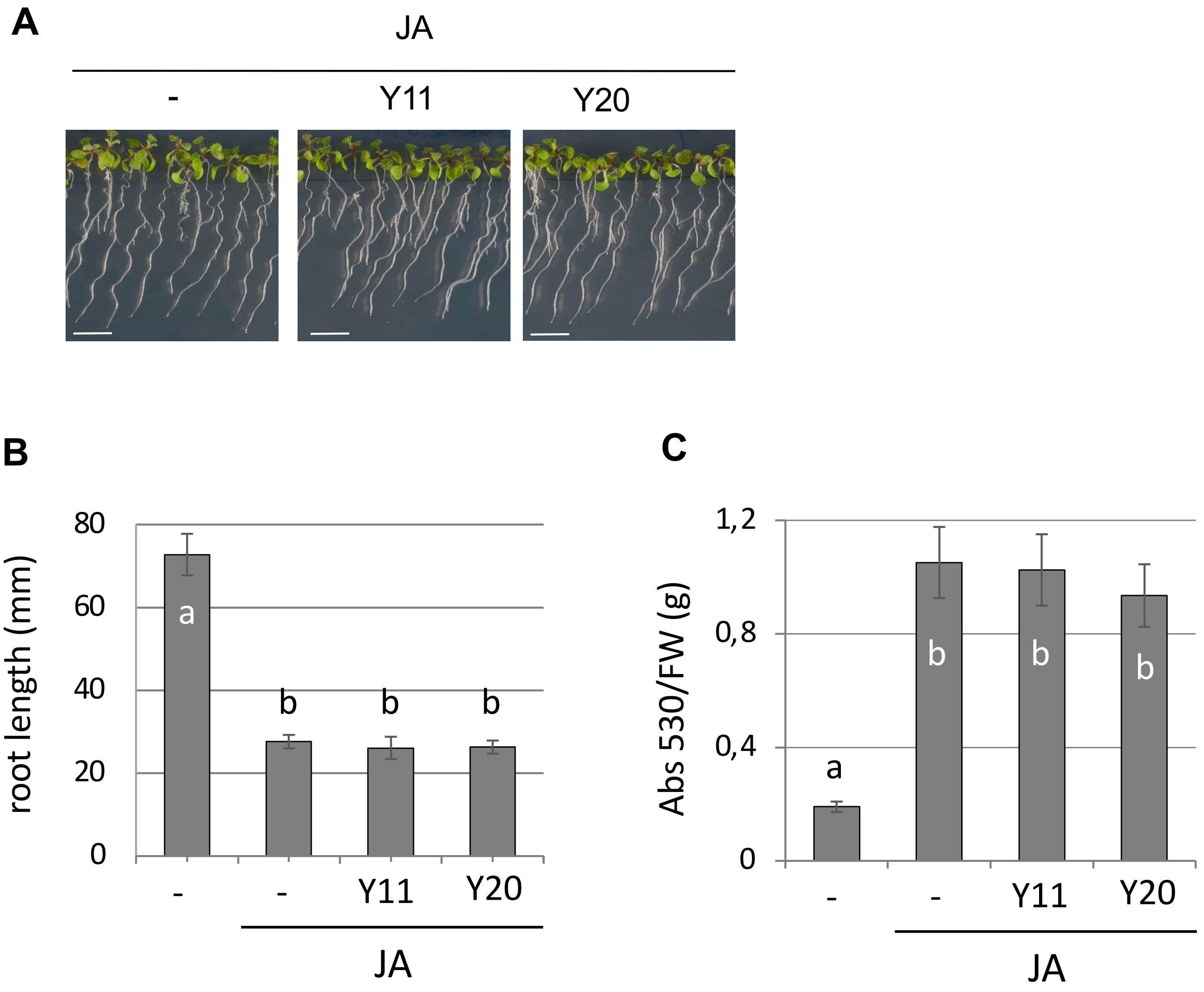


Supplemental Figure S10. **Y11 and Y20 do not prevent JA-mediated responses *in planta***.

(A) 13-day-old WT seedlings germinated in MS media for 3 days and then grown for 10 days in vertical plate in presence of 10 µM JA and 2.5 µM Y11 or Y20. Compounds were dissolved in DMSO, therefore an equivalent volume of DMSO was used as negative control (labelled as -) in. A-C. Lines represent 1 cm.

(B) Root growth inhibition by 15 µM JA of 13-day-old WT seedlings grown for 10 days in vertical plate in presence or absence of 2.5 µM Y11 or Y20. Results are expressed as mean +/- SD of 25-30 plants. One-way ANOVA with post-hoc Tukey HSD Test ($p < 0.01$) analyses define the significant differences.

(C) Anthocyanin accumulation, shown as Absorbance (530) per gram of plant fresh weigh (FW), in 8-day-old WT seedlings grown for 2 days in presence of 50 µM JA with or without 2.5 µM Y11 or Y20. Results are expressed as mean +/- SD. One-way ANOVA with post-hoc Tukey HSD Test ($p < 0.01$) analyses define the significant differences.

A

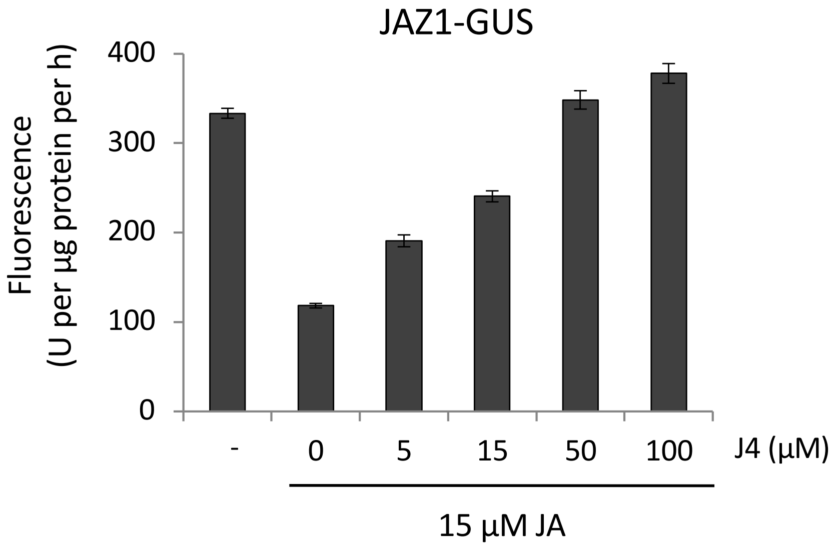


B

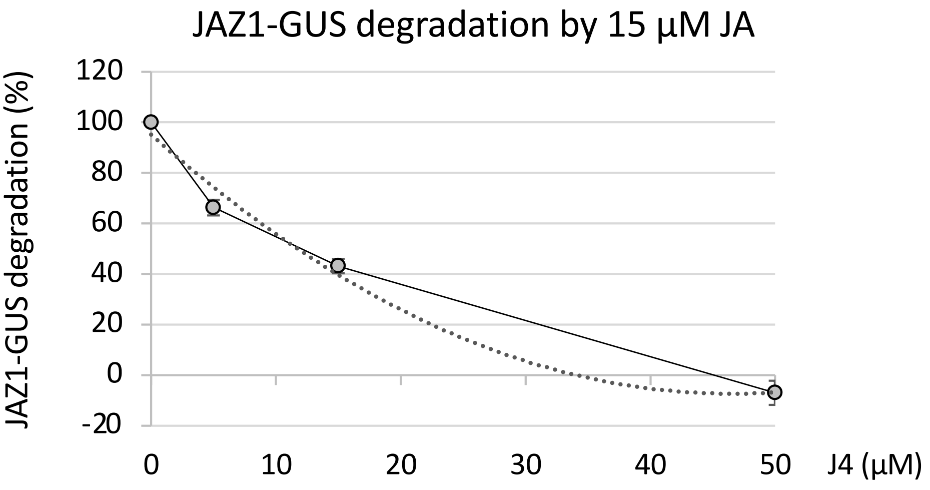


Supplemental Figure S11. **Half maximal inhibitory concentration (IC50) of J4 in JA-promoted JAZ1-GUS degradation.**

(A) Quantification of GUS activity (arbitrary unit U per µg protein per h) in roots of 7-d-old JAZ1-GUS plants. Seedlings (N =20 to 30) were pretreated with the indicated concentrations of J4 for 1 h and then treated with 15 µM JA for 1 hour. Results shown are the mean ± s.d. of seven replicates.

(B) Relative quantification of JAZ1-GUS. Untreated control was set as 100% of GUS activity in roots of 7-d-old JAZ1-GUS plants. Seedlings (N =20 to 30) were pretreated for 1 hour with the indicated concentrations of J4 and then with 15 µM JA for an additional hour. Results shown are the mean ± s.d. of seven replicates. The plain line connects the experimental values, whereas the dotted line is the trendline.

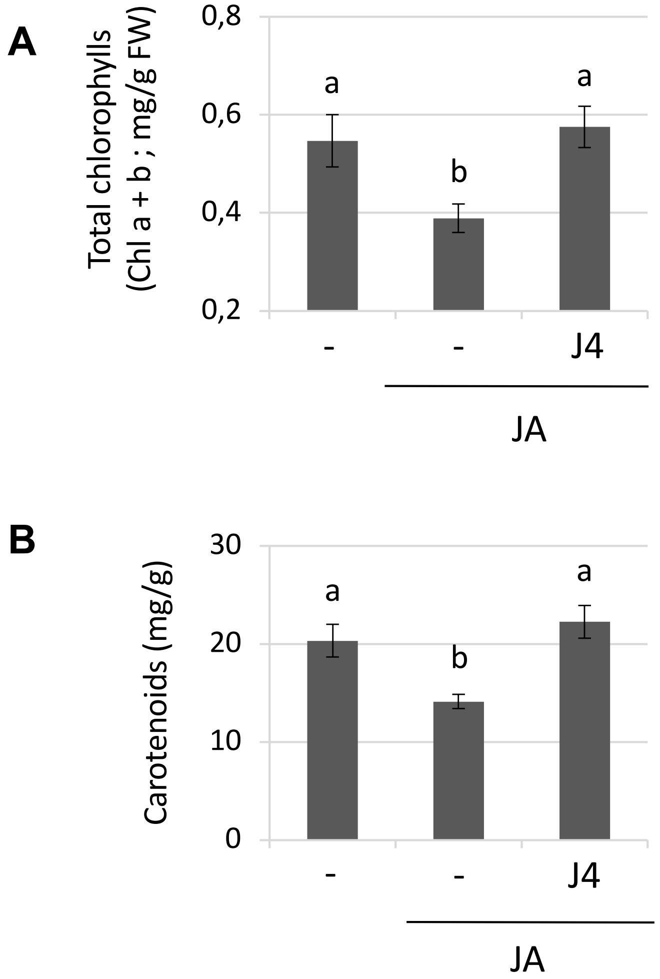


Supplemental Figure S12. **Antagonistic effect of the compound J4 on JA-induced responses in *Nicotiana benthamiana* plants.**

Chlorophyll a and b (A) and carotenoids (B) content of 10-day-old *N. benthamiana* seedlings (N = 7 plants) germinated in vertical plates and then grown for 3 days in presence of 25 µM JA with or without 10 µM J4. The experiments were repeated three times with similar results.

A and B, Bars represent the average value and error bars the standard deviation. One-way ANOVA with post-hoc Tukey HSD Test ($p< 0.01$) analyses define the significant differences.

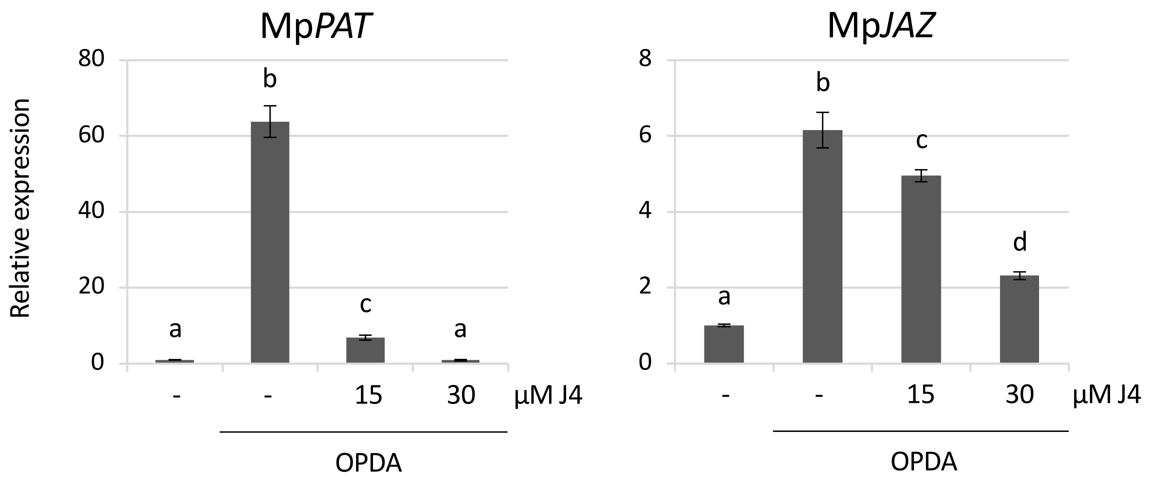


Supplemental Figure S13. **J4 partially inhibits OPDA-mediated transcriptional activation.** Gene expression analysis of Mp*PAT* and Mp*JAZ* in Tak-1 Marchantia plants in response to 10 µM OPDA (oxophytodienoic acid) for 1 hour; plants pre-treated with DMSO (-; untreated control), 15 or 30 µM J4 (+) for 1 hour. Mp*ACT* was used as housekeeping control gene. One-way ANOVA with post-hoc Tukey HSD Test ($p < 0.01$) analyses define the significant differences in gene expression. Each biological sample consisted of tissue pooled from 5-8 plants. Data show mean ± SD of four technical replicates.

Supplemental Table S1. List of all identified molecules, structures, molecular weight, canonical SMILES and commercial ID.

| Compound | srtucture | MW | SMILES | commercial ID (supplaier) | InChI Key |
|---|---|---|---|---|---|
| **J4** | 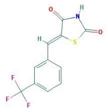 | 273,2 | N1C(SC(\C1=O)=C/c1cc(C(F)(F)F)ccc1)=O | 5721666 (ChemBridge) | NGJLOFCOEOHFKQ-YVMONPNESA-N |
| **Y10** | 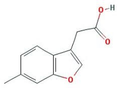 | 189,2 | c1(c2c(cc(cc2)C)oc1)CC(=O)O | 5934888 (ChemBrigde) | ZGVSMROZRSOMIQ-UHFFFAOYSA-N |
| **Y11** | 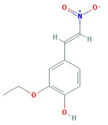 | 209,2 | [N+](\C=C\c1cc(OCC)c(cc1)O)([O-])=O | 6510918 (ChemBrigde) | GAJIQLKTZKVZAJ-AATRIKPKSA-N |
| **Y20** | 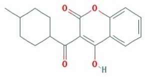 | 286,3 | c1(c(c2ccccc2oc1=O)O)C(C1CCC(CC1)C)=O | 7822375 (ChemBrigde) | KVUQMOJOINIPSW-UHFFFAOYSA-N |
| **J4-L1** | 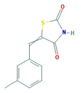 | 219,3 | CC1=CC=CC(=C1)C=C2C(=O)NC(=O)S2 | 5377712 (ChemBrigde) | TURIRIIGEMCNAC-UHFFFAOYSA-N |
| **J4-L2** | 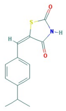 | 247,3 | CC(C)c1ccc(\C=C2\SC(=O)NC2=O)cc1 | 5376783 (ChemBrigde) | GXIDNNRIWNVXTL-YRNVUSSQSA-N |
| **J4-L3** | 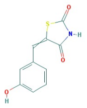 | 221,2 | C1=CC(=CC(=C1)O)C=C2C(=O)NC(=O)S2 | 5378417 (ChemBrigde) | IKLKVFXCODJJRX-UHFFFAOYSA-N |
| **Y17** | 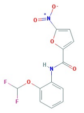 | 297,0 | c1([N+]([O-])=O)oc(C(Nc2c(OC(F)F)cccc2)=O)cc1 | 7758685 (ChemBrigde) | WILYKJXNRSBHHG-UHFFFAOYSA-N |
| **Y18** | 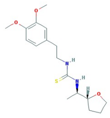 | 336,0 | C(N[C@@H]([C@@H]1OCCC1)C)(=S)NCCc1cc(OC)c(cc1)OC | 7789134 (ChemBrigde) | KILQWPLQZMLGEQ-TZMCWYRMSA-N |
| **J1** | 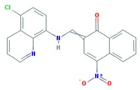 | 377,0 | [N+](c1c2c(cccc2)c(c(c1)\C=N/c1c2c(cccn2)c(cc1)Cl)O)([O-])=O | 5565929 (ChemBrigde) | ZOVIIPWCBYZXMA-UHFFFAOYSA-N |
| **J2** | 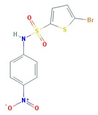 | 363,2 | S(c1sc(Br)cc1)(Nc1ccc([N+]([O-])=O)cc1)(=O)=O | 7598798 (ChemBridge) | XAKPASIXPSBWRB-UHFFFAOYSA-N |
| **J3** | 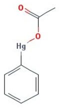 | 336,7 | CC(=O)O[Hg]C1=CC=CC=C1 | LAT003D02 (LACTA) | XEBWQGVWTUSTLN-UHFFFAOYSA-M |
| **J9** | 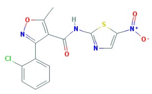 | 364,8 | C1=CC=C(C(=C1)C2=NOC(=C2C(=O)NC3=NC=C(S3)[N](=O)O)C)Cl | 5540228 (ChemBridge) | RKXMPSCTGRFVJB-UHFFFAOYSA-N |
| **J10** | 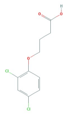 | 249,1 | C1=CC(=C(C=C1Cl)Cl)OCCCC(=O)O | 330041 (Spectrum) | YIVXMZJTEQBPQO-UHFFFAOYSA-N |
| **J11** | 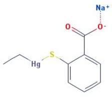 | 404,8 | CC[Hg]SC1=CC=CC=C1C(=O)[O-].[Na+] | 01500572 (Spectrum) | RTKIYNMVFMVABJ-UHFFFAOYSA-L |

Supplemental Table S2

List of primers used for RT-qPCR analyses

| Name | Sequence |
|---|---|
| AtJAZ10 qForw | GAGAAGCGCAAGGAGAGATTAG |
| AtJAZ10 qRev | CTTAGTAGGTAACGTAATCTCC |
| AtOPR3 qFrow | GCATGGAAGCAAGTTGTGGAAGCA |
| AtOPR3 qRev | CATGCGCCCCGTGGATCTCAAT |
| AtTAT3 qFrow | AAGCTGAAGGCCGAGGATGTGTAT |
| AtTAT3 qRev | TCCCGGCCTTGGAAGTAGAATGTT |
| AtACT8 qFrow | CCAGTGGTCGTACAACCGGT |
| AtACT8qRev | TAGTTCTTTTCGATGGAGGAGCTG |
| SlJAZ10 qFrow | GGAACTCACTCTTTCTCCTAGCAAC |
| SlJAZ10 qRev | TGGTGATGAAGGCTCAGACAGCTT |
| SlMYC qFrow | GAGAATTTCAAGGAAGTTCAAGAAT |
| SlMYC qRev | GGCTTTATTTCACACAGAGATAAAA |
| SlPI-II qFrow | GAAAATCGTTAATTTATCCCACCG |
| SlPI-II qRev | ACATACAAACTTTCCATCTTTACCA |
| Sl$\alpha$TUB4 qFrow | AAACAGACCGGCATTTTACAG |
| Sl$\alpha$TUB4 qRev | GGTGATGATGAAGCAGATGGT |
| NbJAZ3 qFrow | CTGAGGCAAAATCTGAACCGGAG |
| NbJAZ3 qRev | GCACCCAATCCAAGCCACAC |
| NbTIFY5 qFrow | GAAAGCACTGAAGAGAAACAAC |
| NbTIFY5 qRev | CCTCCATTCCTCTACTTGCC |
| NbTIFY6b qFrow | AACACAACACCTACTCTCCCC |
| NbTIFY6b qRev | AGAACCGAGCCAATGATGCC |
| Nb$\beta$ACT qFrow | ATGGCAGACGGTGAGGATATTCA |
| Nb$\beta$ACT qRev | GCCTTTGCAATCCACATCTGTTG |
| MpDIR qFrow | CGGAGAAGGTAATTGTCACCACA |
| MpDIR qRev | TCTACCATACAGAGGACGTGATCG |
| MpBHLH4 qFrow | AGAGATTTTGCTCCCTGCAA |
| MpBHLH4 qRev | CATCTGTTGCAAATGCTGCT |
| MpPAT qFrow | GAGATTCACCCCACAAAGAACG |
| MpPAT qRev | GATCTTGGTAACCCTTGAAGTTGG |
| MpJAZ qFrow | ACAGAAGAATGGGTTGTCAGACG |
| MpJAZ qRev | GGACCGGAAGAGACAGTAAGAGTG |
| MpACT qFrow | AGGCATCTGGTATCCACGAG |
| MpACT qRev | ACATGGTCGTTCCTCCAGAC |
